# Multiple conformations facilitate PilT function in the type IV pilus

**DOI:** 10.1101/672212

**Authors:** Matthew McCallum, Samir Benlekbir, Sheryl Nguyen, Stephanie Tammam, John L. Rubinstein, Lori L. Burrows, P. Lynne Howell

## Abstract

Type IV pilus-like systems are protein complexes that polymerize a fibre of pilins. They are critical for virulence in many pathogens. Pilin polymerization and depolymerization are powered by motor PilT-like ATPases thought to possess *C*_2_ symmetry. However, most PilT-like ATPases crystallize with either *C*_3_ or *C*_6_ symmetry and the relevance of these conformations is unclear. Here we determined the X-ray structures of PilT in four unique conformations and used these structures to classify the conformation of available PilT-like ATPase structures. Single particle electron cryomicroscopy (cryoEM) structures of PilT revealed condition-dependent preferences for *C*_2,_ *C*_3_, and *C*_6_ conformations. The physiologic importance of these conformations was validated by co-evolution analysis and functional studies of point mutants, identifying a rare gain-of-function mutation that favours the *C*_2_ conformation. With these data we propose a comprehensive model of PilT function with broad implications for PilT-like ATPases.

## Introduction

Type IV pilus-like (T4P-like) systems are distributed across all phyla of prokaryotic life (1, 2). T4P-like systems include the type IVa pilus (T4aP), type II secretion (T2S) system, type IVb pilus (T4bP), Tad/Flp pilus, Com pilus, and archaellum – with the latter found exclusively in Archaea. These systems enable attachment, biofilm formation, phage adsorption, surface or swimming motility, natural competence, and folded protein secretion in Bacteria and Archaea (3-5), and thus are of vital medical and industrial importance. As many of these systems are critical for virulence in bacterial pathogens (1, 6-8), conserved components could have value as therapeutic targets. Despite the importance of T4P-like systems, basic questions – including how the pilus is assembled and disassembled – remain open.

All systems have at least three conserved and essential elements: a pilus polymer of subunits termed pilins, a pre-pilin peptidase, and a motor (9). The pre-pilin peptidase cleaves the N-terminal leader peptides of pilin subunits at the inner face of the cytoplasmic membrane, leaving the mature pilins embedded in the membrane (10). Polymerization requires extraction of pilins from the membrane by the cytoplasmic motor using energy generated from ATP hydrolysis (11-13). The motor is made up of two well-conserved components: a cytoplasmic ring-like hexameric PilT-like ATPase and a PilC-like inner-membrane platform protein (9). As mature pilins cannot interact directly with the cytoplasmic PilT-like ATPase, their polymerization requires interaction of both the pilins and ATPase with the PilC-like platform protein (9). Cryo-electron tomography (cryoET) studies of the T4aP, T4bP, archaellum, and T2S systems are consistent with localization of the PilC-like protein in the lumen of the hexameric PilT-like ATPase, connecting it to the pilus on the exterior of the inner membrane (14-17). The PilT-like ATPase connects to stator-like components, suggesting that a fixed PilT-like ATPase moves the PilC-like protein (15). Thus, PilT-like ATPases are thought to power pilin polymerization by rotating the PilC-like protein to extract pilins from the membrane and inserting them into the base of the growing pilus polymer (15, 18).

Detailed structural analysis of PilT-like ATPases has advanced our understanding of how the T4P-like motor could insert pilins into the base of a helical pilus. PilT-like ATPase hexamers can be represented as six rigid subunits, termed packing units, held together by flexible linkers (**Figure 1 – figure supplement 1**) (18). Adjacent packing units are oriented in one of two conformations: open (O) or closed (C) (18). PilB is the PilT-like ATPase that powers pilin polymerization in the T4aP. All PilB motor structures determined to date are similar in overall conformation: they exhibit *C*_2_ symmetry with a CCOCCO pattern of O- and C-interfaces around the hexamer (*C*_*2*_^CCOCCO^). In this conformation, the pore of PilB is elongated. Using the heterogeneous distribution of nucleotides in ADP-bound and ADP/ATP-analog-bound PilB crystal structures, we deduced that ATP binding and hydrolysis in the *C*_*2*_^CCOCCO^ PilB structure would propagate conformational changes leading to a clockwise rotation of the elongated pore (18). PilC, bound in the pore, would thus be turned clockwise in 60° increments, while accompanying conformational changes in the PilB subunits would displace PilC out of the plane of the inner membrane, towards the periplasm (18). If a pilin is inserted at each clockwise increment these motions would build a one-start, right-handed helical pilus (18), consistent with cryoEM structures (19, 20).

T4aP polymers can be rapidly depolymerized at the base, resulting in fibre retraction. PilT is the PilT-like ATPase that powers T4aP depolymerisation (21). We applied the same analysis used to deduce the movements in PilB to the *C*_*2*_ symmetric structure of PilT from *Aquifex aeolicus* (PilT^Aa^, PDB 2GSZ (22)) (18). We found that this protein had an OOCOOC pattern of interfaces (*C*_*2*_^OOCOOC^), which would give the impression of counter-clockwise rotation of the elongated pore and potential downward movement of PilC (18). Thus, we proposed that PilT may act like PilB in reverse, consistent with powering pilus depolymerization (18). This analysis highlighted the importance of clarifying the symmetry and pattern of O- and C-interfaces in PilT-like ATPases when interpreting PilT-like ATPase structures and defining their mechanisms.

In contrast to PilB structures, which exhibit only *C*_2_ symmetry, PilT has been crystallized in a variety of conformations with *C*_6_ symmetry (22-24). Other PilT-like ATPases have crystallized in conformations with *C*_2_, *C*_3_, and *C*_6_ symmetries (12, 25, 26). Multiple potential crystallographic conformations of PilT and PilT-like ATPases suggest that the *C*_*2*_^OOCOOC^ conformation may not represent the active PilT retraction motor (22, 27). Further, the *C*_*2*_^OOCOOC^ PilT structure was determined using a homolog from the Aquificae (22) and may not reflect a conformation typical of PilT from Proteobacteria, where most of the phenotypic analyses of the T4aP have been conducted. The specific conformation adopted by the PilT motor, and the details of retraction remain to be clarified.

Here we crystallized PilT in four unique conformations, including the highest resolution structure to date of a hexameric PilT-like ATPase. These structures allowed the identification of conserved O- and C-interface contact points that were used to differentiate the conformations of available PilT-like ATPase structures into six unique classes. To examine the conformations adopted by PilT and PilB in a non-crystalline state, we determined their structures using cryoEM. CryoEM structures of PilT revealed a clear preference for *C*_2_ and *C*_3_ conformations in the absence of nucleotide or with ADP, and the *C*_6_ conformation in the presence of ATP. These structures were validated by co-evolution analysis and functional analysis of point mutants. A gain-of-function mutation with increased *in vivo* activity was identified, and its cryoEM analysis revealed a preference for the *C*_2_ conformation. From these data, we propose a comprehensive model of PilT function with broad implications for all PilT-like ATPases.

## RESULTS

### PilT crystallizes in *C*_*3*_ and pseudo-*C*_*3*_ symmetric conformations

To gauge the reproducibility of previously crystallized PilT conformations and to assess if additional conformations could be observed, PilT4 from *Geobacter metallireducens* (PilT^Gm^) was crystallized. PilT^Gm^ was selected because we previously crystallized PilB from *G. metallireducens* to derive models for PilB-mediated extension (18). There are four PilT orthologs in *Geobacter*, and PilT4 is the primary retraction ATPase in *Geobacter sulfurreducens* (28). In the absence of added nucleotide, PilT^Gm^ only crystallized after reductive methylation. These crystals diffracted to 3.3 Å, and the structure, determined by molecular replacement, revealed a *C*^3^ symmetric hexamer in the asymmetric unit (**Figure 1A**). No nucleotide was found in this structure, although density consistent with sulfate, present in the crystallization conditions, was observed in the nucleotide-binding site (**Figure 1B**). By comparison with the interfaces in PilB^Gm^, we categorized the interfaces between the packing units of PilT^Gm^ as alternating between O and C: OCOCOC. This *C*_*3*_^OCOCOC^ conformation has not previously been observed for PilT.

**Figure 1.**
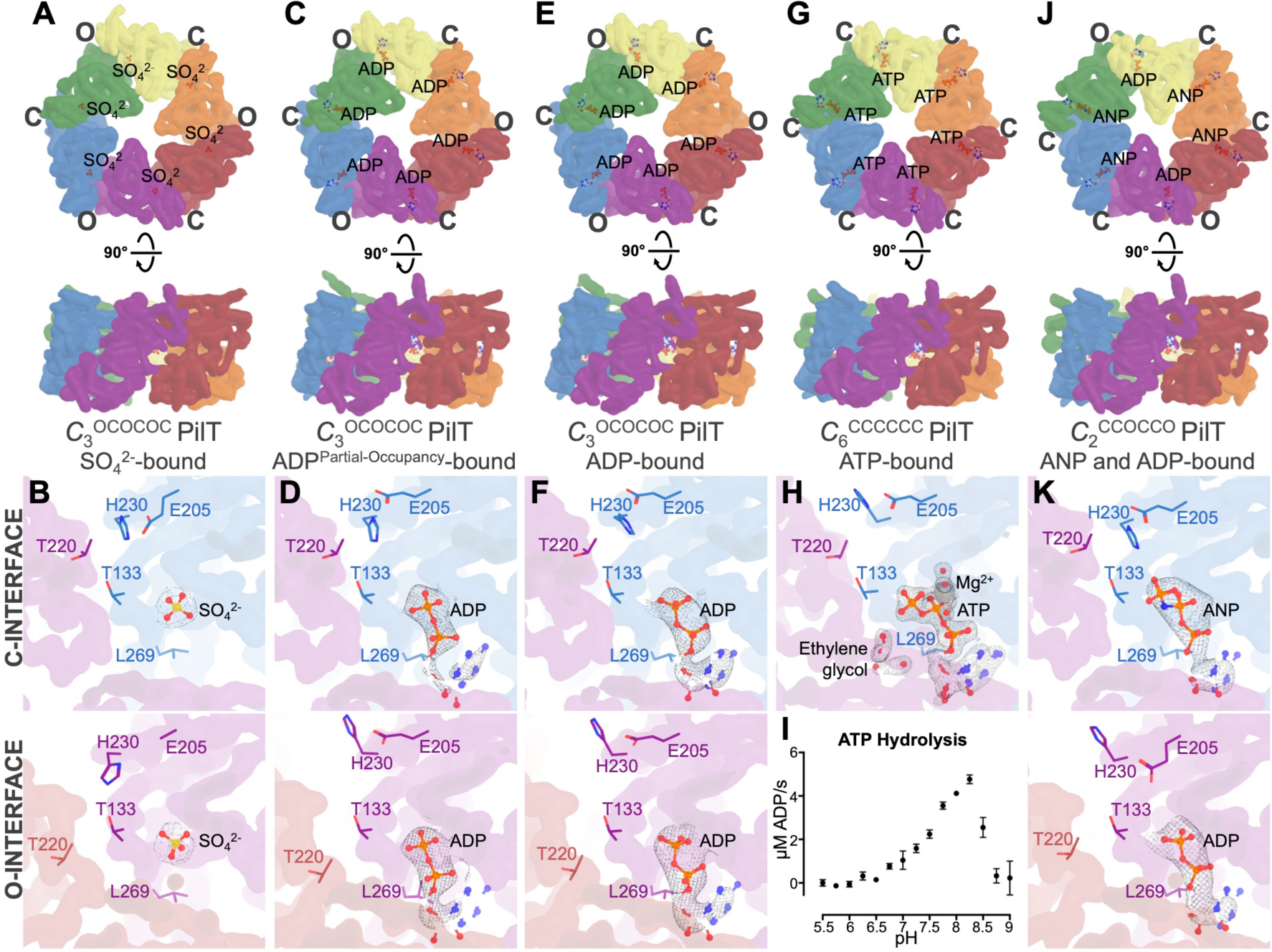
PilT from *G. metallireducens* (PilT^Gm^) crystallizes in multiple conformations. Individual packing units (N2D^n^ and CTD^n+1^) are uniquely coloured. A smoothed transparent surface representation of the main chain is shown. The ligand bound in the nucleotide-binding site is shown in stick representation. **B, D, F, H**, and **J**, a cross-section of the C- and O- interfaces are shown on the top and bottom panels, respectively. Select side chains discussed in the text are shown for reference. 2F_o_-F_c_ maps shown with *I/σ(I)* levels as indicated below. **A-B**, Crystal structure of methylated PilT^Gm^ with an alternating pattern of O- and C- interfaces – in the *C*_3_^OCOCOC^ conformation. Nucleotide-binding sites of representative O- and C- interfaces in the methylated PilT structure are shown; *I*/ *σ*(*I)* of map is 2. **C-D**, crystal structure of PilT^Gm^ without added nucleotide with an alternating pattern of O- and C- interfaces – in the *C*_3_^OCOCOC^ conformation. Nucleotide-binding sites of representative O- and C- interface are shown; In keeping with the partial occupancy of the nucleotide the *I*/*σ*(*I)* of map is 1.5. **E-F**, crystal structure of PilT^Gm^ washed with ATP prior to crystallization in the *C*_3_^OCOCOC^ conformation. Nucleotide-binding sites of representative O- and C- interfaces; *I*/*σ*(*I)* of map is 2. **G-H**, crystal structure of PilT^Gm^ incubated with ATP during crystallization at pH 6.5 in the *C*_6_^CCCCCC^ conformation. Nucleotide-binding sites of a representative C- interface in the structure is shown; *I*/*σ*(*I)* of map is 2. **I**, *In vitro* ATP hydrolysis by PilT^Gm^ from pH 5.5 to 9.0. **J-K**, crystal structure of PilT^Gm^ incubated with ANP during crystallization with a CCOCCO pattern of O- and C- interfaces – in the *C*_2_^CCOCCO^ conformation. Nucleotide-binding sites of representative O- and C- interfaces in the structure are shown; *I*/*σ I* of map is 2.

We hypothesized that this conformation in the absence of nucleotide could have resulted from the methylation process, rather than representing a physiologically-relevant PilT conformation. Therefore, to crystallize PilT^Gm^ in the absence of added nucleotide without reductive methylation, we extensively optimized the buffer (see Experimental Procedures). The resulting crystals diffracted to 3.0 Å, and the structure was solved by molecular replacement with the entire hexamer present in the asymmetric unit (**Figure 1C**). This structure was approximately *C*_*3*_ symmetric with alternating O- and C-interfaces, similar to the methylated *C*_*3*_^OCOCOC^ PilT structure. However, small deviations make this structure pseudo-*C*_*3*_ symmetric. Despite not adding nucleotide, density consistent with ADP allowed nucleotide with partial occupancy to be modeled in all nucleotide-binding sites (**Figure 1D**). These nucleotides were likely carried over from *Escherichia coli* during protein purification. Nucleotide was absent in the methylated PilT^Gm^ structure, possibly due to the lengthy methylation protocol or competition with sulfate during crystallization.

An isomorphous structure with high ADP occupancy was obtained by pre-incubating PilT^Gm^ with Mg^2+^ and ATP, then removing unbound nucleotide prior to crystallization (**Figure 1E and 1F**). This structure is consistent with the *C*_3_^OCOCOC^ structure reflecting a post-hydrolysis ADP-bound conformation. The isomorphic low-occupancy ADP and high-occupancy ADP *C*_*3*_^OCOCOC^ PilT structures have an RMSD^C*α*^ of 0.6 Å per hexamer. The RMSD^C*α*^ of these two structures with the methylated *C*_*3*_^OCOCOC^ PilT structure is 1.9 Å per hexamer.

### PilT^Gm^ also crystallizes in a *C*_*6*_ symmetric conformation

Modifying the protocol so that exogenous ATP was not removed prior to crystallization yielded distinct PilT^Gm^ crystals that diffracted to 1.9 Å, the highest resolution to date for any hexameric PilT-like ATPase family member. The structure was solved by molecular replacement with three protomers in the asymmetric unit. In the crystal, two nearly identical *C*_*6*_ symmetric hexamers could be identified (**Figure 1G**). Compared with the interfaces of PilB^Gm^, all six interfaces in this hexamer are closed. This conformation is denoted *C*_6_^CCCCCC^.

Density consistent with Mg^2+^ and surprisingly for an active ATPase, ATP, could be modeled in the active sites (**Figure 1H**). The ribose moiety of ATP puckers in two alternate conformations consistent with the small number of direct protein contacts to the O2’ of ATP (**Figure 1J**) observed. These conformations are consistent with C2’ exo and C2’ endo low-energy ATP ribose conformations (29). In addition, two ethylene glycol molecules from the cryoprotectant solution could be modelled in every packing unit interface. The ethylene glycol was introduced after the crystals formed and bound to Arg-83 and Arg-278, next to the nucleotide-binding site. As these crystals formed only when the pH was less than or equal to 6.5, we hypothesised that PilT^Gm^ may not have ATPase activity at acidic pH. When we assayed PilT^Gm^ ATPase activity over a broad-range of pH values, we could not measure ATPase activity below pH 6.5 (**Figure 1I**). H230 of the HIS box motif in PilT is predicted by Rosetta (30) to have a pKa of 6.5; thus H230 deprotonation might be important for efficient ATP hydrolysis. The corresponding histidine in PilB coordinates the nucleotide γ-phosphate, but in the *C*_*6*_^CCCCCC^ PilT structure, H230 is facing away from the γ-phosphate in each of the nucleotide-binding sites. We propose that the protonation state of H230 affects its preferred rotamer conformation and subsequently the catalytic activity of PilT. Since H230 is conserved in all PilT-like ATPases (18), this pH dependency may be conserved in other PilT-like ATPases.

### PilT^Gm^ also crystallizes in a PilB-like *C*_*2*_ symmetric conformation

To determine what conformation PilT^Gm^ adopts above pH 6.5 with an ATP analog, we also crystallized PilT^Gm^ with Mg^2+^ and a non-hydrolysable ATP analogue, ANP (adenylyl-imidodiphosphate or AMP-PNP) at pH 8. Unlike previous PilT^Gm^ crystals that formed after 16 h and were stable for weeks, these crystals took a week to form and stayed crystalline for two days before dissolving. These crystals diffracted anisotropically to 4.1, 6.7, and 4.0 Å resolution along *a*\*, *b*\*, and *c*\* reciprocal lattice vectors, respectively. The structure was solved by molecular replacement, and a hexamer was present in the asymmetric unit (**Figure 1J**). Despite the low resolution, density consistent with ANP could be modeled into four of the six nucleotide-binding sites (**Figure 1K**). The density in the other two sites was consistent with ADP. It is possible, given the slow and transient crystallization, that ANP partially hydrolyzed, yielding a transient ADP/ANP mixture that facilitated formation of these particular crystals. In this case, decay to ADP is likely non-catalytic. This structure of PilT^Gm^ is *C*_2_ symmetric but distinct from that of the *C*_*2*_^OOCOOC^ PilT^Aa^ structure (22) (RMSD^C*α*^ 6.8 Å / hexamer). Surprisingly, the pattern of O- and C- interfaces between packing units is PilB-like: CCOCCO, or *C*_*2*_^CCOCCO^.

### ATP but not ADP binding correlates with C-interfaces

As PilT^Gm^ crystallized in multiple conformations and the resolution and quality of the electron density was sufficient to resolve the bound nucleotide, we looked for correlations between ADP or ATP/ATP-analog binding and the O- or C-interfaces. In the ADP-bound *C*_*3*_^OCOCOC^ PilT^Gm^ structures, the estimated occupancy of ADP was similar in the O- and C- interfaces (**Figure 1D** and **1F**). In the *C*_*6*_^CCCCCC^ PilT^Gm^ structure, all interfaces bound to ATP (**Figure 1J**). In the *C*_*2*_^CCOCCO^ PilT^Gm^ structure, the four C-interfaces are bound to the ATP analog ANP, while the O-interfaces appear to be bound to ADP (**Figure 1H**). Thus, there is a correlation between bound ATP (or ATP-analog) and closed interfaces, suggesting that ATP facilitates closure of interfaces in PilT^Gm^. In contrast, there is no correlation between bound ADP and O- and C-interfaces, suggesting that both O- and C-interfaces in PilT may have a similar affinity for ADP, and ADP may be insufficient to induce or maintain interface closure. This scenario contrasts with PilB^Gm^ structures, where ADP is correlated with C-interfaces (18).

### Conserved interactions facilitate O- and C-interface formation in PilT

We established previously that in the C-interface of PilB^Gm^, T411 contacts H420, and in the O-interface, R455 contacts T411 (18). To comprehensively define the interfaces in PilT^Gm^, we graphically plotted the O- and C-interface contacts of the PilT^Gm^ crystal structures, as well as previously published PilT structures (**Figure 2**). For this analysis, a contact was defined as any atom (main-chain or side-chain) of a residue within 4 Å of any atom from another residue. To compensate for imperfect rotamers in low-resolution structures, the definition of contacts for this analysis was expanded to also include main-chain atoms in one residue within 8 Å of a main-chain atom from another residue.

**Figure 2.**
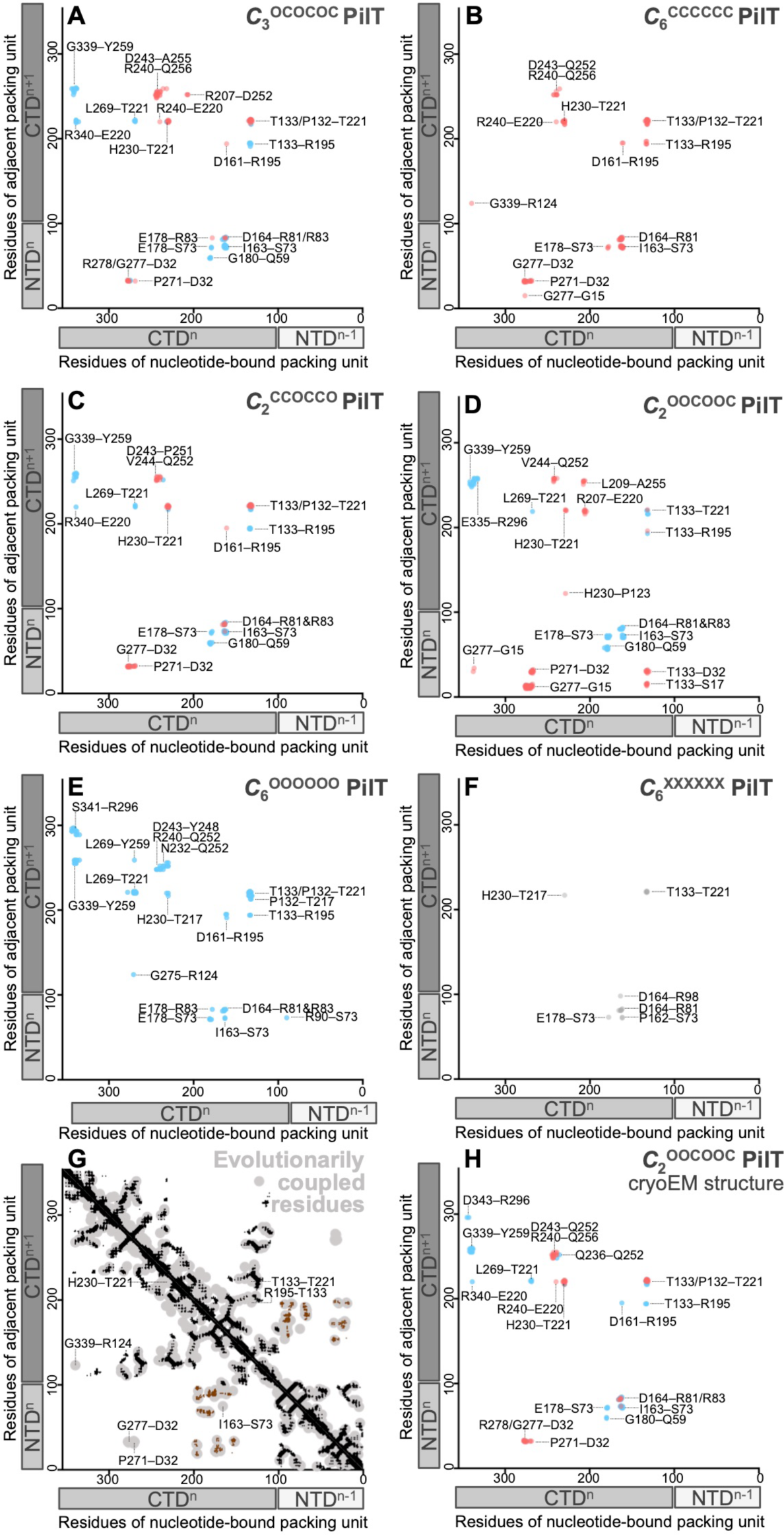
Contact maps for packing unit interfaces reveal similarities between distinct PilT conformations. This analysis indicates whether packing unit interface residues are in proximity in crystal structures (**A-F**), based on evolutionarily coupled residues (**G**), and in a cryoEM structure (**H**). Inter- and intra-chain contacts between packing units colour coded based on C- (red) or O- (blue) interface; the X interface is shown in grey. For reference, the linear domain architecture of individual packing units is shown as box cartoons beside the axes with each chain shaded distinctly. If nucleotide were bound between two adjacent packing units, the ‘nucleotide-bound packing unit’ would be bound to the nucleotide by the Walker A motif. The ‘adjacent packing unit’ is the adjacent packing unit in the interface. PDB coordinates representative of particular conformations were used for this analysis: PilT^Gm^ crystal structures presented herein were used for the *C*_3_^OCOCOC^ (**A**, PDB 6OJY), *C*_6_^CCCCCC^ (**B**, PDB 6OJX), and *C*_2_^CCOCCO^ (**C**, PDB 6OKV) conformations. PilT^Aa^ crystal structures were used for the *C*_2_^OOCOOC^ (**D**, PDB 2GSZ) and *C*_6_^XXXXXX^conformations (**F**, PDB 2EWV). The PilT^Gs^ crystal structure was used for the *C*_6_^OOOOOO^ conformation (**E**, PDB 5ZFQ). To compare with the *C*_2_^OOCOOC^ crystal structure, the *C*_2_^OOCOOC^ conformation PilT^Gm^ cryoEM structure presented herein is used (**H**, PDB 6OLL). To minimize confusion when comparing between species – which have different residue identities and numbering than PilT^Gm^ – the residue labels represent the corresponding residue in PilT^Gm^. For clusters of residues that are in proximity, only the most prominent contacts (salt-bridges and dipolar interactions) are labeled. **G**, evolutionarily coupled residues in PilT (light grey) are compared with the structure of PilT^Gm^. Tertiary structure contacts (not including intra-chain interactions between packing units) are noted in black. N2D^n^ to CTD^n+1^ contacts (that create individual packing units) are noted in brown. Evolutionarily coupled residues could ambiguously correspond to residue contacts that start from the nucleotide-bound packing unit or start from the adjacent packing unit. Both possibilities are presented, thus there is mirror symmetry across the x=y plane; the tertiary structure and N2D^n^ to CTD^n+1^ contacts are presented likewise. O- or C- interface contacts identified in **A-F** are labeled if they overlap with evolutionarily conserved residue pairs that are not clearly accounted for by the tertiary structure or N2D^n^ to CTD^n+1^ contacts.

Despite disparate hexameric conformations, the PilT structures have fairly consistent O- and C-interface contacts. Residues in the C-interface that contact one another include **P271–D32**, G277–D32, **H230–T221**, T221–T133, **R195–D161**, and D164–R81; bolded contacts are specific to the C-interface. Residues in the O-interface that consistently contact one another include **G339–Y259, G180–Q59**, R195–T133, T221–T133, E178–S73, I163–S73, and D164–R81 and bolded contacts are specific to the O-interface. Many of these residues are part of the conserved HIS (T221 through H230) and ASP (E160 through E164) box motifs (31, 32). Involvement in the O- and C-interfaces explains why several residues in these motifs are conserved, even though only H230 and E164 contact ATP or magnesium, respectively (18, 22). The contacts observed in the O- and C-interfaces are similar to those found in the PilB structure: T221 contacts H230 in the C-interface (T411 and H420 in PilB^Gm^), while T221 contacts L269 (T411 and R455 in PilB^Gm^) in the O-interface.

Based on this information, C- and O-interfaces can be easily distinguished by measuring the Cα distance between inter-chain residues T221 to H230 and T221 to L269. In the C-interfaces of PilT^Gm^ structures, the mean Cα distances for T221–H230 and T221–L269 are 5.7 ± 0.3 Å (mean ± SD) and 13.7 ± 0.8 Å, respectively. In the O-interfaces of PilT^Gm^ structures, the mean Cα distances for T221–H230 and T221–L269 are 12.7 ± 1 Å and 8.6 ± 0.4 Å, respectively. The T221–H230 and T221–L269 distances are closer in the C- and O-interfaces, respectively, permitting easy interface classification.

In subsequent analyses, we used an arbitrary cut-off of < 9 Å for the T221–H230 distance and >12 Å for T221–L269 to define the C-interface. To define the O-interface, we used an arbitrary cut-off of >12 Å for T221–H230 and < 11 Å for T221–L269. T221–H230 and T221– L269 distances were measured across representative PilT-like ATPase structures, to identify their C- and O-interfaces (**Table 1**). The mean T221–H230 and T221–L269 backbone distances in PilT-like ATPase structures are, respectively, 6 ± 1 Å and 16 ± 3 Å in C-interfaces, and 16 ± 4 Å and 9 ± 1 Å in O-interfaces.

**Table 1.**
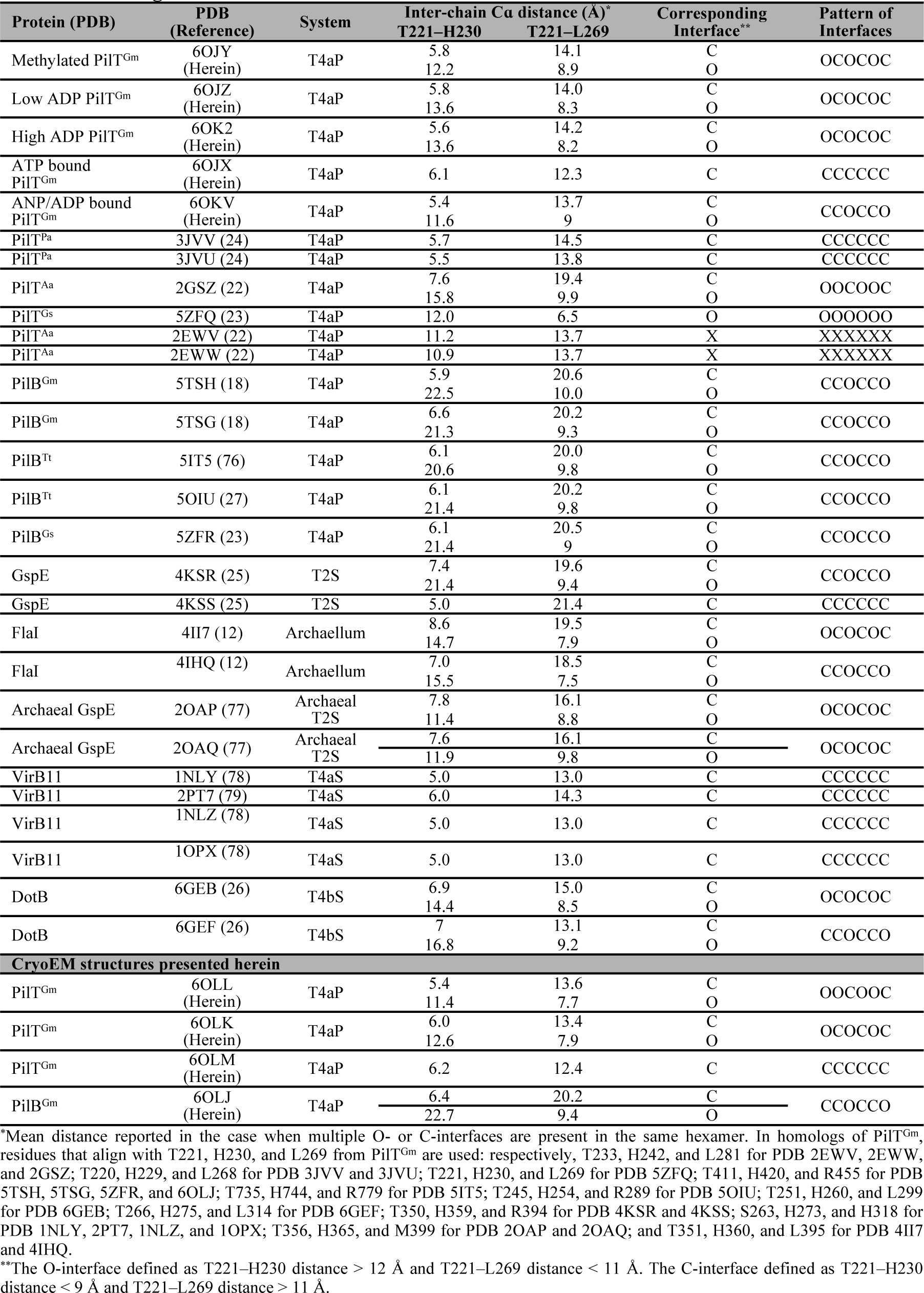
Defining the O- and C-interfaces across PilT-like ATPase structures.

### Structures of PilT-like ATPases can be subdivided into six conformational states

Characterization of the O- and C-interfaces in all PilT-like ATPase structures allowed most to be placed into one of five different states based on their conformation: *C*_*2*_^CCOCCO^, *C*_*3*_^OCOCOC^, *C*_*6*_^CCCCCC^, *C*_*6*_^OOOOOO^, and *C*_*2*_^OOCOOC^ (**Figure 3**). These states reflect all possible arrangements of O- and C-interfaces that maintain rotational symmetry. A GspE structure (PDB 4KSS) and all VirB11 crystal structures (PDB 2PT7, 2GZA, 1NLY, 1NLZ, and 1OPX) are *C*_*6*_^CCCCCC^ (**Figure 3A**). In addition to the ATP-bound PilT^Gm^ structure described herein, the PilT structures from *Pseudomonas aeruginosa* (PilT^Pa^) are also in the *C*_*6*_^CCCCCC^ conformation (PDB 3JVV and 3JVU) (**Figure 3A**). The three *C*_*3*_^OCOCOC^ PilT structures described here are the only examples of this conformation class determined to date (**Figure 3B**). A FlaI and a DotB structure (PDB 4II7 and 6GEB, respectively), as well as two Archaeal GspE2 structures (20AP and 2OAQ) have *C*_*3*_^OCOCOC^ conformations (**Figure 3B**). Other GspE, FlaI, and DotB structures (PDB 4KSR, 4IHQ, and 6GEF) fall into the *C*_*2*_^CCOCCO^ state (**Figure 3C**). All available PilB structures are *C*_*2*_^CCOCCO^ (**Figure 3C**). Our *C*_*2*_^CCOCCO^ PilT^Gm^ structure is the only example to date in this state (**Figure 3C**). PilT^Aa^ is the only example of the *C*_*2*_^OOCOOC^ conformational state (PDB 2GSZ) (**Figure 3D**). Similarly, only PilT4 from *Geobacter sulfurreducens* (PilT^Gs^) exhibits a *C*_*6*_^OOOOOO^ conformation (**Figure 3E**). This classification scheme suggests that PilT and PilT-like ATPase crystal structures have a high fidelity for O- or C-interfaces and rotational symmetry.

**Figure 3.**
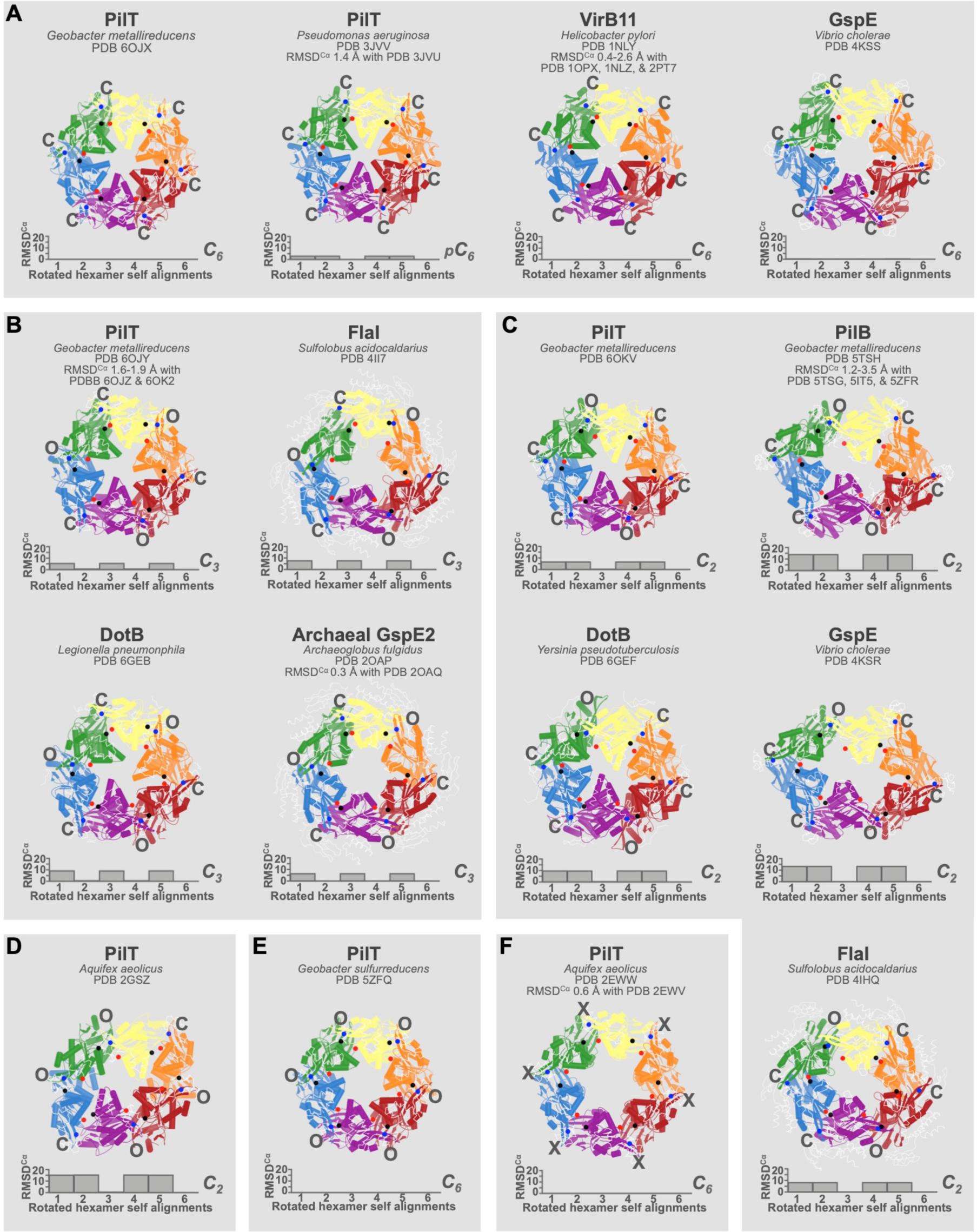
All PilT-like ATPase structures can be divided into six unique conformations. Structures are shown as cartoons with individual packing units (N2D^n^ plus CTD^n+1^) uniquely coloured. Structural elements – like domains, α-helices, or *β*-sheets – that are not well conserved across all PilT-like ATPases are shown as thin white ribbons to highlight similarities between conformations. The hexamers are labelled with their protein name, followed by the species of origin, followed by their PDB identifier, and finally the PDB identifiers of similar structures with the range of RMSD^Cα^ (over the full hexamer) of these structures aligned with the shown structure. **Black spheres**, the α-carbons of the residues that align with T221 from PilT^Gm^. **Red spheres**, the α-carbons of the residues that align with H230 from PilT^Gm^. **Blue spheres**, the α-carbons of the residues that align with L269 from PilT^Gm^. The interface between packing units is annotated as O, C, or X, as determined using the T221, H230, and L269 distances in Table 1. Below each cartoon hexamer is a **graph** of the RMSD^Cα^ of each hexamer aligned with the same hexamer offset by *n* chains (see Methods); the *x*-axis is the value *n*. The symmetry is noted to the right of each graph. **A**, PilT-like ATPase conformations with *C*_6_ or pseudo-*C*_6_ (p*C*_6_) symmetry and six C- interfaces, i.e. the *C*_6_^CCCCCC^ conformation. **B**, PilT-like ATPase conformations with *C*_3_ symmetry or pseudo-*C*_3_ (p*C*_3_) symmetry and alternating O- and C-interfaces, i.e. the *C*_3_^OCOCOC^ conformation. **C**, PilT-like ATPase conformations with *C*_2_ symmetry and the CCOCCO pattern of interfaces, i.e. the *C*_2_^OOCOOC^ conformation. **D**, the only PilT-like ATPase structure with *C*_2_ symmetry and the OOCOOC pattern of interfaces, i.e. the *C*_2_^OOCOOC^ conformation. **E**, the only PilT-like ATPase structure with *C*_6_ symmetry and six O- interfaces, i.e. the *C*_6_^OOOOOO^ conformation. **F**, PilT-like ATPase conformations with *C*_6_ symmetry, and six X-interfaces, i.e. the *C*_6_^XXXXXX^ conformation.

Cross-referencing PilT structural classification with packing-unit interface contacts (**Figure 2**) revealed that there are contacts unique to some conformational states. For example, the G339–R124 and G275–R124 contacts are specific to the *C*_*6*_^CCCCCC^ and *C*_*6*_^OOOOOO^ PilT conformational states, respectively (**Figure 2B & E**). There are many contacts unique to the *C*_*2*_^OOCOOC^ PilT structure (**Figure 2D**), though this may be a result of the evolutionary distance of PilT^Aa^ from Proteobacteria. There are no contacts unique to the *C*_*2*_^CCOCCO^ and *C*_*3*_^OCOCOC^ PilT^Gm^ structures (**Figure 2A and 2C**).

PilT^Aa^ was crystallized in a distinct *C*_*6*_ symmetric conformation bound to ATP (PDB 2EWW) or ADP (PDB 2EWV) (**Figure 3F**). The two structures can be considered isomorphic (RMSD_^C^_*α* 0.6 Å / hexamer) and are distinct from other PilT-like ATPase structures. There is almost no interface between packing units in these structures, so the packing units appear to be held in place by the crystal contacts suggesting this conformation may be uncommon outside of a crystal lattice. The distances of T221–H230 and T221–L269 are both atypically large (11 and 14 Å, respectively) (**Table 1**). These distances suggest that the packing units have neither a C- nor O-interface and thus we refer to it as an X-interface. Thus, these *C*_6_ symmetric PilT^Aa^ structures represent a distinct conformational state: *C*_*6*_^XXXXXX^. The H242–R229 salt-bridge is a major constituent of the X interface; these residues are H230 and T217, respectively, in PilT^Gm^ (**Figure 2F**). As R229 in PilT^Aa^ is not conserved in PilT^Gm^, the X-interface or *C*_6_^XXXXXX^ conformation may not be critical to function. Supporting this hypothesis, it was suggested that these *C*_*6*_^XXXXXX^ PilT^Aa^ structures are not in a conformation that could facilitate ATP hydrolysis (22).

### Residues in the interface between packing units are evolutionarily coupled

One explanation for the perceived heterogeneity in PilT and PilT-like ATPases is that these proteins are conformationally heterogenous and that the crystallization process selects for just one of many possible conformations. To probe for this possibility, we used the EVcouplings server (33, 34) to identify residues that co-evolve in PilT (**Figure 2G**). Residues that contact one another – and are needed for biological function – tend to be evolutionarily coupled, and therefore this analysis can be used to independently validate structural analyses (33, 34).

The evolutionarily-coupled residues of PilT were consistent with its tertiary structure contacts, as well as the contacts that support the packing unit. There were also evolutionarily coupled residues clustered around G277–D32, P271–D32, H230–T221, T221–T133, R195–T133, and G339–R124 consistent with the C-interface, and I163–S73 consistent with the O- or C-interface. This analysis unambiguously demonstrates the phylogenetic conservation of the C- interface across PilT orthologs. The G339–R124 contact is specific to the *C*_*6*_^CCCCCC^ PilT^Gm^ structure specifically validating the biological relevance of this conformation.

The quality of sequence alignments is worse near the C terminus of PilT and thus the power to find evolutionarily conserved residues is lower near the C terminus. Also, some O- and C-interface contacts overlap graphically with tertiary or packing-unit forming contacts, obscuring their identification in this type of analysis. Thus, while this analysis validates the *C*_*6*_^CCCCCC^ PilT structure and C-interface, other non-crystallographic techniques are required to test the biological relevance of particular conformations.

### 2D cryoEM analysis of PilT^Gm^ reveals conformational heterogeneity

To examine what conformation(s) PilT can adopt in a non-crystalline environment we used cryoEM. We found that PilT^Gm^ adopted preferred orientations on the EM grid with its symmetry axis normal to the air-water interface (top-views), preventing calculation of 3D maps from untilted images. Nonetheless, comparison of 2D class average images with projections of PilT crystal structures (**Figure 4A**) allowed for assignment of most 2D class averages into one of the six defined PilT conformational states. In the absence of nucleotide, the 2D class averages of PilT^Gm^ were a mixture of the *C*_*3*_^OCOCOC^ and *C*_2_^OOCOOC^ conformational classes in a 45:55 ratio (**Figure 4B**).

**Figure 4.**
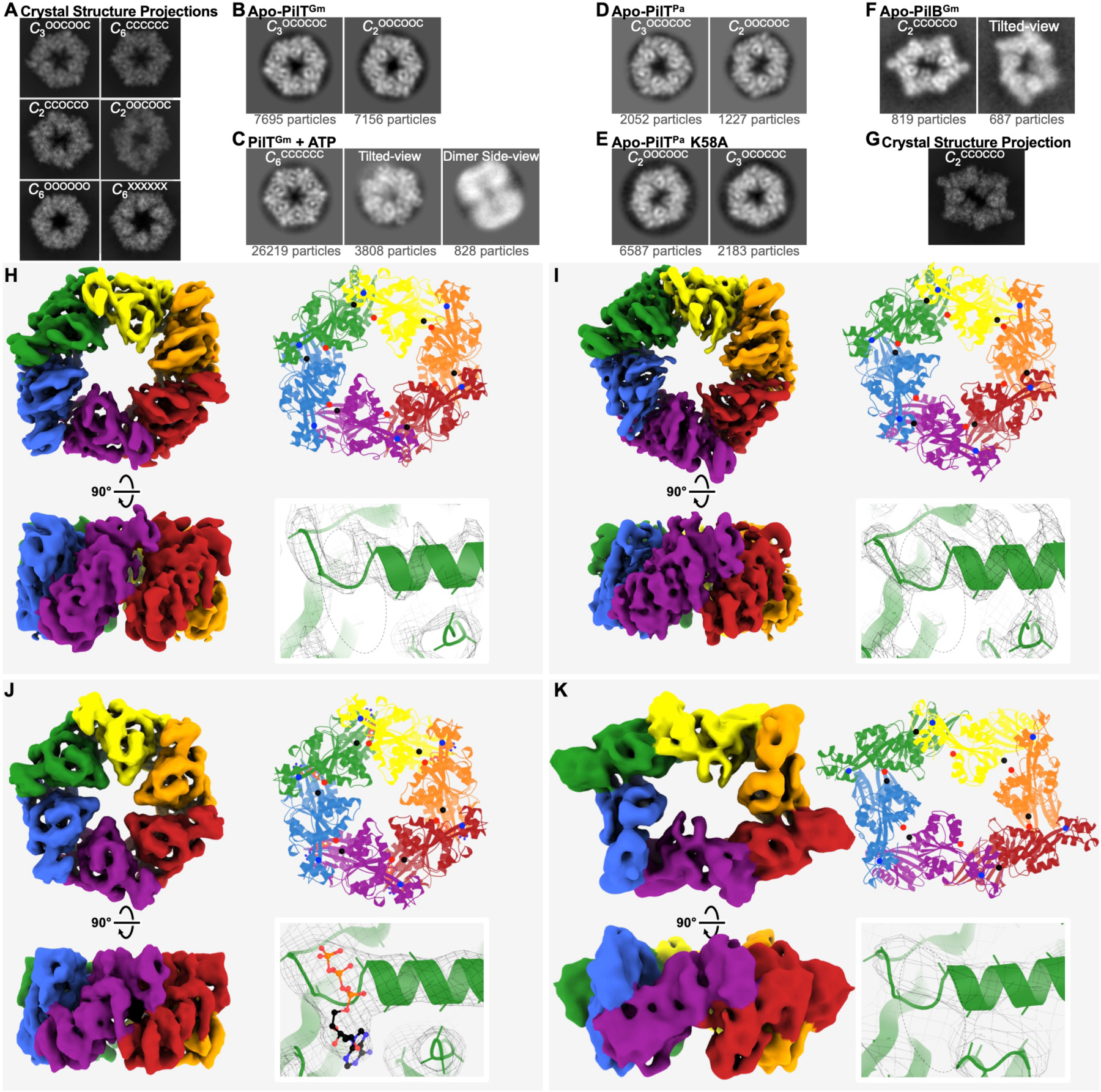
CryoEM structure analysis reveals the conformation preferences of PilT and PilB. **A**, PilT crystal structures were used to simulate top-view cryoEM projections. For this analysis, hexamers representative of *C*_3_^OCOCOC^, *C*_6_^CCCCCC^, *C*_2_^CCOCCO^, *C*_2_^OOCOOC^, *C*_6_^OOOOOO^, and *C*_6_^XXXXXX^ conformations were extracted from the corresponding PDB coordinates: 6OJY (published herein), 6OJX (published herein), 6OKV (published herein), 2GSZ, 5ZFQ, and 2EWV, respectively. Using these simulated projections the 2D class averages in **B-E** are annotated with their predicted conformation (white font). For scale, the grey squares shown in **B-F** are 16 nm wide relative to the class averages. **B**, Images of representative 2D class averages from apo-PilT^Gm^ particles. **C**, Images of representative 2D class averages from particles of PilT^Gm^ pre-incubated with 1 mM ATP. **D**, Images of representative 2D class averages from apo-PilT^Pa^ particles. **E**, Images of representative 2D class averages from apo-PilT^Pa^ particles with the K58A mutation. **F**, Images of representative 2D class averages from apo-PilB^Gm^ particles. **G**, the hexamer from a PilB^Gm^ crystal structure (5TSG) was extracted and used to simulate a cryoEM projection. **H-K**, Sharpened cryoEM maps coloured as in Figure 1 by packing units (left); the top-view of models built into these maps (cartoons, top-right) with spheres shown for the α-carbons of residues that align with T221, H230, and L269 as in Figure 3 (black, red, and blue spheres, respectively); and a zoomed view of the Walker A motif (bottom-right) showing the quality of the sharpened maps (mesh) and presence or absence of nucleotide. A dotted outlined oval shows the location of the nucleotide-binding site. **H**, CryoEM map of PilT in the *C*_2_^OOCOOC^ conformation to 4.1 Å resolution determined in the absence of added nucleotide. **I**, CryoEM map of PilT in the *C*_3_^OCOCOC^ conformation to 4.0 Å resolution also determined in the absence of added nucleotide **J,** CryoEM map of PilT pre-incubated with 1 mM ATP in the *C*_6_^CCCCCC^ conformation to 4.4 Å resolution. **K**, CryoEM map of PilB in the *C*_2_^CCOCCO^ conformation to 7.8 Å resolution determined in the absence of added nucleotide.

In the presence of 0.1 mM ADP and 1 mM ADP, the ratio of the particles in the *C*_*3*_^OCOCOC^ and *C*_2_^OOCOOC^ states shifted from 45:55 to 54:46 and 71:29, respectively (**Figure 4 – figure supplement 1**). Thus, adding ADP favours the *C*_*3*_^OCOCOC^ conformation, consistent with crystallographic *C*_*3*_^OCOCOC^ structures of PilT^Gm^ with ADP bound in every interface. There may be an unoccupied nucleotide-binding site available in the *C*_2_^OOCOOC^ conformation that upon binding ADP converts to the *C*_*3*_^OCOCOC^ conformation. This hypothetical unoccupied nucleotide-binding site might prefer to bind a different nucleotide such as ATP, since mM quantities of ADP were required to favour the *C*_*3*_^OCOCOC^ conformation.

### Purified PilT from *P. aeruginosa* also demonstrates conformation heterogeneity

PilT^Pa^ has been crystallized in the *C*_*6*_^CCCCCC^ conformation (**Figure 3A**) both in the absence of nucleotide and in the presence of an ATP analog (24). We hypothesized that purified PilT^Pa^ may also exhibit the *C*_*3*_^OCOCOC^ or *C*_*2*_^OOCOOC^ conformation in the absence of nucleotide in a non-crystalline environment. Thus, cryoEM analysis was performed with purified PilT^Pa^. Analysis of the 2D class averages were – like PilT^Gm^ – consistent with the *C*_*3*_^OCOCOC^ and *C*_*2*_^OOCOOC^ conformations (**Figure 4D**). The particle distribution ratio between *C*_3_^OCOCOC^ and *C*_2_^OOCOOC^ class averages was 63:37.

### 3D cryoEM analysis of PilT^Gm^ validates coexisting *C*_2_^OOCOOC^ and *C*_3_^OCOCOC^ conformations

To confirm our interpretation of the 2D class averages, we obtained sufficient views of the hexamers to calculated 3D maps by tilting the specimen (35). CryoEM specimens of PilT^Gm^ without nucleotide were tilted by 40° during data collection. Using heterogeneous refinement in *cryoSPARC* (36) and per-particle determination of contrast transfer function parameters (37), two distinct maps could be obtained from the data without enforcing symmetry: one ∼*C*_3_ map and one ∼*C*_2_ map, both at ∼4.4 Å resolution. The particle distribution ratio was 48:52 between *C*_3_ and *C*_2_ maps, respectively; similar to that obtained by 2D classification. Applying their respective symmetries during refinement yielded 4.0 Å and 4.1 Å resolution maps, respectively (**Figure 4H and 4I**).

Molecular models could be built into these maps by fitting and refining rigid packing units of the PilT^Gm^ crystal structures (**Figure 4H and 4I**). Assessment of local resolution suggests that the non-surface-exposed portions of the map are at higher-resolution than the rest of the complex (**Figure 4 – figure supplement 2**). No density consistent with nucleotide was identified in these structures, presumably due to the low resolution of the maps. The model built into the *C*_*3*_ symmetric map is consistent with the *C*_*3*_^OCOCOC^ PilT structure (RMSD^C*α*^ 1.3 Å / hexamer). Before symmetry was applied to this map, it more closely matched the methylated *C*_3_ symmetric structure of PilT than the pseudo-*C*_3_ symmetric structures, suggesting that the slight asymmetry of the latter is a crystallographic artefact. The model built into the *C*_*2*_ symmetric map was not consistent with any PilT^Gm^ crystal structure. Annotation of its packing unit interfaces revealed that it has a *C*_*2*_^OOCOOC^ conformation, consistent with the PilT^Aa^ crystal structure (**Table 1**). Thus, the cryoEM structures confirm that the *C*_*2*_^OOCOOC^ and *C*_*3*_^OCOCOC^ conformations observed for PilT^Aa^ and PilT^Gm^, respectively, were not crystal artefacts. Further, these maps suggest that available crystal structures have over-simplified our view of PilT-like ATPases as they do not capture the multiple stable conformations accessible in a given condition.

While the *C*_*2*_^OOCOOC^ PilT^Gm^ cryoEM structure validates the *C*_*2*_^OOCOOC^ PilT^Aa^ crystal structure, the two are distinct (RMSD^C*α*^ of 6.4 Å / hexamer). Analyzing the packing unit interfaces of the *C*_2_^OOCOOC^ PilT^Gm^ cryoEM structure reveals that they are nearly identical to the interfaces in the PilT^Gm^ *C*_*2*_^CCOCCO^ and *C*_*3*_^OCOCOC^ conformations (**Figure 2H**).

### CryoEM analysis of PilT^Gm^ in the presence of ATP consistent with *C*_6_^CCCCCC^ conformation

Since PilT hydrolyzes ATP slowly and cryoEM samples can be frozen within minutes of sample preparation, we opted to determine the conformation of PilT^Gm^ incubated briefly with ATP. In these conditions, the top-view 2D class averages of PilT^Gm^ corresponded only to the *C*_*6*_^CCCCCC^ conformational class, consistent with the ATP-bound *C*_*6*_^CCCCCC^ PilT crystal structure (**Figure 4C**). A small minority of 2D class averages appeared to be tilted- or stacked side-views, permitting 3D map construction. Only one map with ∼*C*_6_ symmetry could be constructed, and applying *C*_6_ symmetry during refinement resulted in a 4.4 Å resolution map (**Figure 4J**). The molecular model built from this map is consistent with the *C*_*6*_^CCCCCC^ crystal structure (RMSD^C*α*^ 0.6 Å) and density in the nucleotide-binding sites is consistent with ATP (**Figure 4J**).

In an attempt to reproduce the conditions that we postulated led to the *C*_*2*_^CCOCCO^ PilT crystal structure, PilT^Gm^ was incubated with mixtures of ATP and ADP. In the presence of 1 mM ATP and ADP, or 1 mM ATP and 0.1 mM ADP, only class averages consistent with the *C*_*6*_^CCCCCC^ conformation could be identified (**Figure 4 – figure supplement 1**). This analysis does not support the reproducibility of the *C*_*2*_^CCOCCO^ PilT conformation in a non-crystalline environment, nor the *C*_*6*_^OOOOOO^ or *C*_*6*_^XXXXXX^ PilT conformations, which were not identified in any condition. The cryoEM experiments suggest that in the absence of its protein-binding partners *in vitro*, at approximately physiological ATP and ADP concentrations, PilT^Gm^ is predominantly found in the *C*_*6*_^CCCCCC^ conformation.

### CryoEM analysis of PilB^Gm^ consistent with *C*_2_^CCOCCO^ conformation

During the course of our studies, an 8 Å cryoEM structure of PilB from *T. thermophilis* (PilB^Tt^) was published that revealed the motor domains of PilB^Tt^ adopt the *C*_*2*_^CCOCCO^ conformation in a non-crystalline environment (27). No conformational heterogeneity was reported (27). To determine whether this homogeneity was observed in a Proteobacteria PilB, we performed cryoEM analysis of PilB from *G. metallireducens* (PilB^Gm^) in the absence of nucleotide. Projection of the PilB *C*_2_^CCOCCO^ crystal structure revealed that the PilB^Gm^ top-view 2D class averages were consistent with the *C*_2_^CCOCCO^ conformation (**Figure 4F and 4G**). A 3D map was calculated at ∼7.8 Å resolution (**Figure 4K**) and the model built into this map is also consistent with the *C*_2_^CCOCCO^ PilB structure (RMSD^C*α*^ 2.3 Å / hexamer, PDB 5TSG). Thus, cryoEM analysis reveals that PilB preferentially adopts the *C*_2_^CCOCCO^ conformation in multiple species. This is in contrast to PilT^Gm^ and PilT^Pa^ which both adopt *C*_2_^OOCOOC^ and *C*_3_^OCOCOC^ conformations in similar conditions. These results suggest that the preferred conformation(s) are conserved within – but not between – PilT-like ATPase sub-families, consistent with distinct conformation preferences facilitating PilB-like or PilT-like functions.

### The *C*_6_^CCCCCC^ and the *C*_2_^OOCOOC^ or *C*_3_^OCOCOC^ containing conformations are essential for PilT retraction

The cryoEM structures of PilT^Gm^ and PilB^Gm^ were determined in the absence of other components of the T4aP system. To explore the functional importance of these PilT conformations *in vivo*, we introduced mutations targeting the packing unit interface into *P. aeruginosa,* a model organism for studying PilT function. From our analysis of key contacts (**Figure 2**), we mutated residues that should alter packing unit interfaces and thus overall conformations. As controls, we mutated the catalytic glutamate (E204A) and the γ-phosphate coordinating HIS-box histidine (H229A), which eliminate twitching motility in *P. aeruginosa* (11). E204A and H229A mutants lost twitching motility and accumulated extracellular PilA, while the E204A also led to PO4 phage resistance, consistent with a retraction defect (**Figure 5A**).

**Figure 5.**
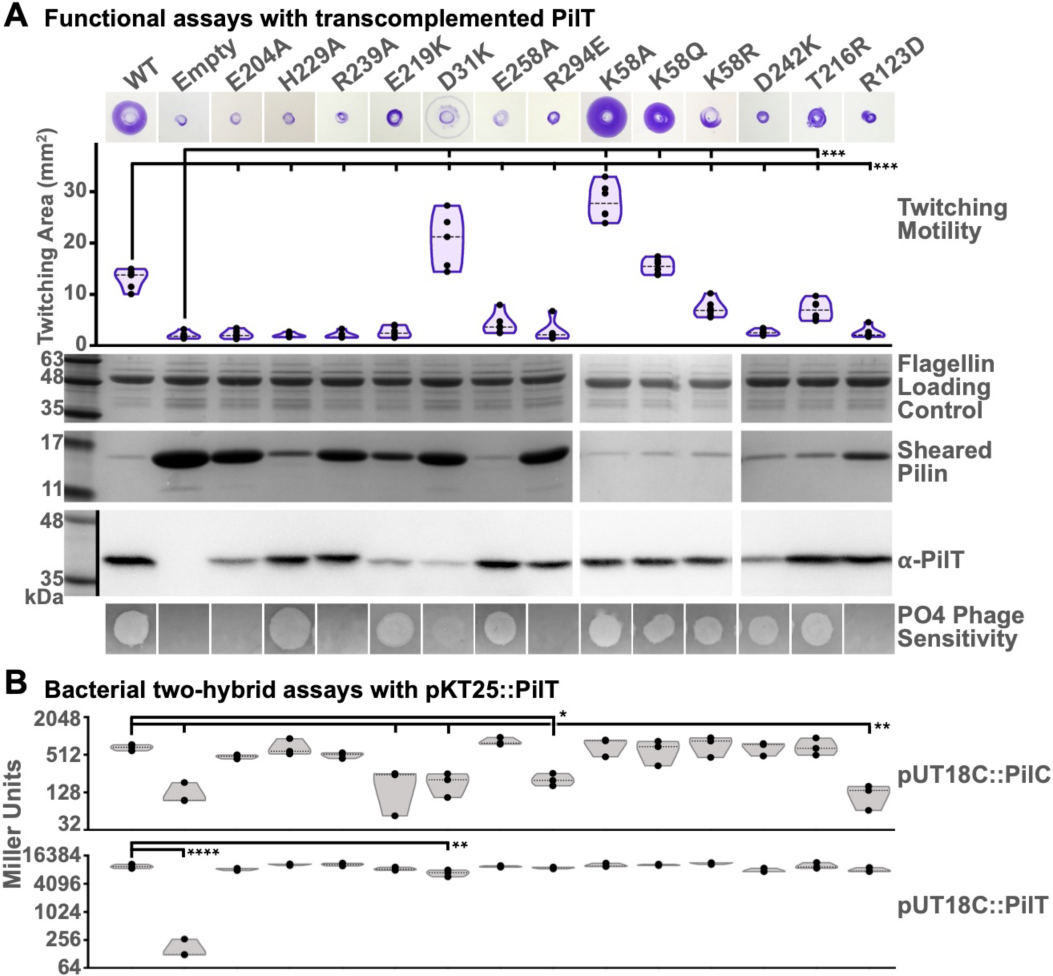
Multiple conformations and both O- and C-interfaces are essential for PilT *in vivo* function. **A**, A *P. aeruginosa* PAO1 PilT deletion mutant was trans-complemented with wildtype (WT), a vector control (Empty), and various site-mutants as indicated. Twitching motility was assayed by crystal violet staining (purple zones) and the areas covered by the bacteria were measured (purple violin plot). Mean twitching areas were compared with the one-way ANOVA test, noted with black lines; reported p-values were less than 0.001 (^***^). Trans-complemented bacteria were grown on a surface; pilins and flagellins were sheared from the surface of the bacteria and assayed for by SDS-PAGE analysis, while whole cell lysates were western-blotted with an anti-PilT antibody (middle). PO4 phage was spotted onto double-layer agar inoculated with the trans-complemented bacterial; transilluminated images of the resulting zones of lysis are shown (bottom). **B**, Quantitative bacterial two-hybrid analysis of PilT^Pa^ mutants. The mutations in **A** were introduced into the pKT25::PilT^Pa^ two-hybrid vector. These were then co-transformed with either pUT18C::PilC^Pa^ (top grey violin plot) or pKT25::PilT^Pa^ (bottom grey violin plot) in the adenylate cyclase mutant, *E. coli* BTH101 and assayed for interaction by *β*-galactosidase activity measured in Miller Units. The mean Miller units were compared with the one-way ANOVA test, noted with black lines; reported p-values were less than 0.02 (^*^), 0.01 (^**^), or 0.0001 (^****^).

In PilT^Gm^ structures, residue R240 participates in the C-interface, E220, D32 and D243 are at both O- and C-interfaces, and F259, R296 and Q59 are at the O-interface (**Figure 2**). In PilT^Pa^ these correspond to R239, E219, D31, E258, D242, R294, and K58, respectively. Mutation of most of these residues abrogated twitching motility, increased extracellular PilA, and led to PO4 phage resistance (**Figure 5A**). These data suggest that the *C*_6_^XXXXXX^ PilT conformation, lacking O- or C-interfaces, is insufficient for PilT function. Likewise, these experiments imply that neither the *C*_6_^CCCCCC^ or *C*_6_^OOOOOO^ PilT conformations – lacking O- or C-interfaces, respectively – are sufficient for PilT function. The E219K and D31K mutants had reduced stability, complicating their interpretation (**Figure 5A**). The side chains of these residues participate in the C-interface but not the O-interface. Curiously, despite the instability of D31K and corresponding accumulation of PilA, D31K had significantly increased twitching motility and was partially susceptible to PO4 phage infection (**Figure 5A**). One interpretation of this phenotype is that PilT misfolded in most bacteria expressing the D31K mutant leading to accumulation of extracellular PilA, but in a subpopulation of bacteria the D31K mutant protein folded properly and unexpectedly facilitated increased twitching motility. Alternatively, it is possible that the D31K mutation reduced binding of the antibody used to detect PilT. The K58Q mutant in PilT^Pa^ had wild-type twitching motility, extracellular pilin accumulation, and were sensitive to PO4 phage, consistent with this residue being glutamine in wild-type PilT^Gm^. Surprisingly, the K58A mutant had 2-fold increased twitching motility and decreased levels of extracellular PilA indicative of hyper-retraction, while the conservative K58R mutation had reduced twitching motility (**Figure 5A**).

Mutations were also introduced at residues that are important for particular conformations. The R124 residue in PilT^Gm^ stabilizes the *C*_6_^CCCCCC^ and *C*_*6*_^OOOOOO^ conformations, as it forms a salt bridge with the backbone carbonyl of G339 and G275, respectively (**Figure 2**). R124 in PilT^Gm^ aligns with R123 in PilT^Pa^. The R123D mutation in PilT^Pa^ eliminated twitching motility, prevented phage infection, and accumulated extracellular pilins, consistent with a retraction defect (**Figure 5A**). This result suggests that either the *C*_6_^CCCCCC^ or *C*_*6*_^OOOOOO^ PilT conformations is essential for retraction. In the *C*_*6*_^OOOOOO^ conformation T217 forms a polar interaction with H230 in the (**Figure 2**) so its mutation to arginine would be expected to eliminate that conformation. T217 in PilT^Gm^ aligns with T216 in PilT^Pa^. The T216R mutant in PilT^Pa^ had slightly decreased twitching motility, and wild-type pilin accumulation and phage infection (**Figure 5A**), suggesting that *C*_*6*_^OOOOOO^ conformation does not play a major role in PilT retraction. These results imply that the *C*_6_^CCCCCC^ conformation is necessary for PilT function. In combination with the mutations targeting O- and C-interfaces (above), these results indicate that the *C*_6_^CCCCCC^ conformation and at least one O- and C-interface containing conformation are essential for PilT retraction. This O- and C-interface containing conformation is likely *C*_2_^OOCOOC^ or *C*_3_^OCOCOC^ as they were the only PilT conformations with O- and C-interfaces reproduced by cryoEM analysis.

### Hyper-twitching K58A mutant favours *C*_2_^OOCOOC^ conformation

Based on the corresponding residues in the O-interface of PilT^Gm^, K58 is predicted to be in proximity of residues H179 and R180 in the O-interface of PilT^Pa^. This could cause electrostatic repulsion to the O-interface. Thus, the K58A mutation may stabilize the O-interface and favour conformations with more O-interfaces. To test this hypothesis, cryoEM analysis was performed on purified K58A PilTPa. The ratio of the particles in the *C*_*3*_^OCOCOC^ and *C*_2_^OOCOOC^ classes shifted from 63:37 for wild-type PilT^Pa^ (**Figure 4D**) to 25:75 for the K58A mutant (**Figure 4E**). This result is consistent with the K58A mutation stabilizing the O-interface and thus favouring the *C*_2_^OOCOOC^ conformation that has more O-interfaces than the *C*_3_^OCOCOC^ conformation.

### The *C*_6_^CCCCCC^ conformation and an O-interface containing conformation are important for binding PilC

We hypothesised that the observed retraction defects reflected the inability of some PilT mutants to adopt a conformation compatible with PilC binding. To test this hypothesis, we used bacterial-two hybrid analysis (BACTH) to quantify the interaction between PilT mutants and PilC (**Figure 5B**). BACTH has been used previously to demonstrate an interaction between PilT and PilC (38). Each PilT mutant was capable of homomeric interactions, consistent with proper protein folding in this assay, with the exception of the D31K mutant, consistent with its putative stability defect (**Figure 5B**). The O-interface-targeting R294E mutant and *C*_*6*_^CCCCCC^-targeting R123D mutant also had reduced PilC interactions (**Figure 5B**). These results indicate that the *C*_*6*_^CCCCCC^ conformation and an O-interface containing conformation are important for binding PilC.

## Discussion

Herein we demonstrate the conformational heterogeneity of PilT. This protein can be crystallized in conformations consistent with all PilT-like ATPase conformational states defined here. We present several novel PilT^Gm^ crystal structures and demonstrate that multiple conformations of PilT^Gm^ and PilT^Pa^ coexist in solution. We show that specific PilT conformations are important for *in vivo* function and interaction with PilC. This study builds unifying concepts and terminology that enable a better understanding of PilT-like ATPase function.

We found that the inter-chain distances between T221–H230 and T221–L269 can be used to easily and quantitatively define O- and C-interfaces. This simple definition enables the conformational state of the hexamer to be defined and would easily be missed if only individual chains are annotated. Given the conservation of these inter-chain distances, we predict that these residues have functional significance. The catalytic glutamate is thought to abstract a proton from a water molecule for subsequent hydroxyl nucleophilic attack of the γ-phosphate of ATP (18). Given the location of H230 adjacent to the catalytic glutamate, H230 might then abstract this proton and shuttle it to T221 in the C-interface. From T221 the proton could be passed directly or indirectly via T133 to the recently hydrolyzed inorganic phosphate. If this were the case, involving T221 from an adjacent packing unit in proton shuttling would prevent efficient ATP hydrolysis in the O-interface prior to its closure. Consistent with this proposed mechanism, we found that PilT^Gm^ ATPase activity is pH sensitive in the range consistent with histidine protonation, and that in PilT^Gm^ structures H230 faces away from the nucleotide-binding site in most O-interfaces and towards the nucleotide-binding site in most C-interfaces.

Our highest resolution crystal structure of a hexameric PilT-like ATPase determined to date also revealed new features. The ATP ribose moiety can adopt multiple conformations due to the lack of interactions with its O2’ hydroxyl. It is thus not surprising that ATP-analogs with fluorophores attached at the O2’ position have been shown to bind PilT-like ATPases (39, 40). We also found two ethylene glycol molecules in the packing unit interface, suggesting that small molecules could target this interface. Targeting the nucleotide-binding site or packing unit interface to inhibit ATPase activity in T4P-like systems may have therapeutic value as these systems are a major virulence factor for many pathogens. Indeed, there are two recent reports of small molecule inhibitors of the *Neisseria meningitidis* T4aP that target pilus assembly and disassembly dynamics to reduce virulence (41, 42). One of these drugs directly targets PilB (41), although whether this drug binds the nucleotide binding site or packing unit interface is not yet clear.

Our cryoEM maps establish the coexistence of both *C*_*2*_^OOCOOC^ and *C*_*3*_^OCOCOC^ conformational classes in PilT in the absence of nucleotide or with added ADP. We also identified the *C*_6_^CCCCCC^ conformation in the presence of ATP or approximately physiological concentrations of ATP and ADP. Based on these analyses we propose a model explaining how the conformation of PilT – and probably other PilT-like ATPases – changes *in vitro* depending on the nucleotides present (**Figure 6B**). We also show by cryoEM analysis, in accordance with the recently determined PilB^Tt^ structure (27), that PilB^Gm^ adopts the *C*_*2*_^CCOCCO^ conformation in a non-crystalline state. Although the structures of these proteins were determined in isolation, the conformations observed are likely biologically relevant, as PilB and PilT are not always engaged with PilC or the T4aP. No core T4aP proteins are unstable in the absence of PilB or PilT (43), cryoET analysis of the T4aP in *M. xanthus* shows a significant portion of T4aP systems without attached PilT-like ATPases (14), and PilB and PilT migrate dynamically within some bacteria while the core T4aP proteins are anchored in the cell envelope (44, 45). Thus, we propose that at physiological ATP and ADP concentrations, when it is not engaged with the T4aP, PilT preferentially adopts the *C*_*6*_^CCCCCC^ conformation.

**Figure 6.**
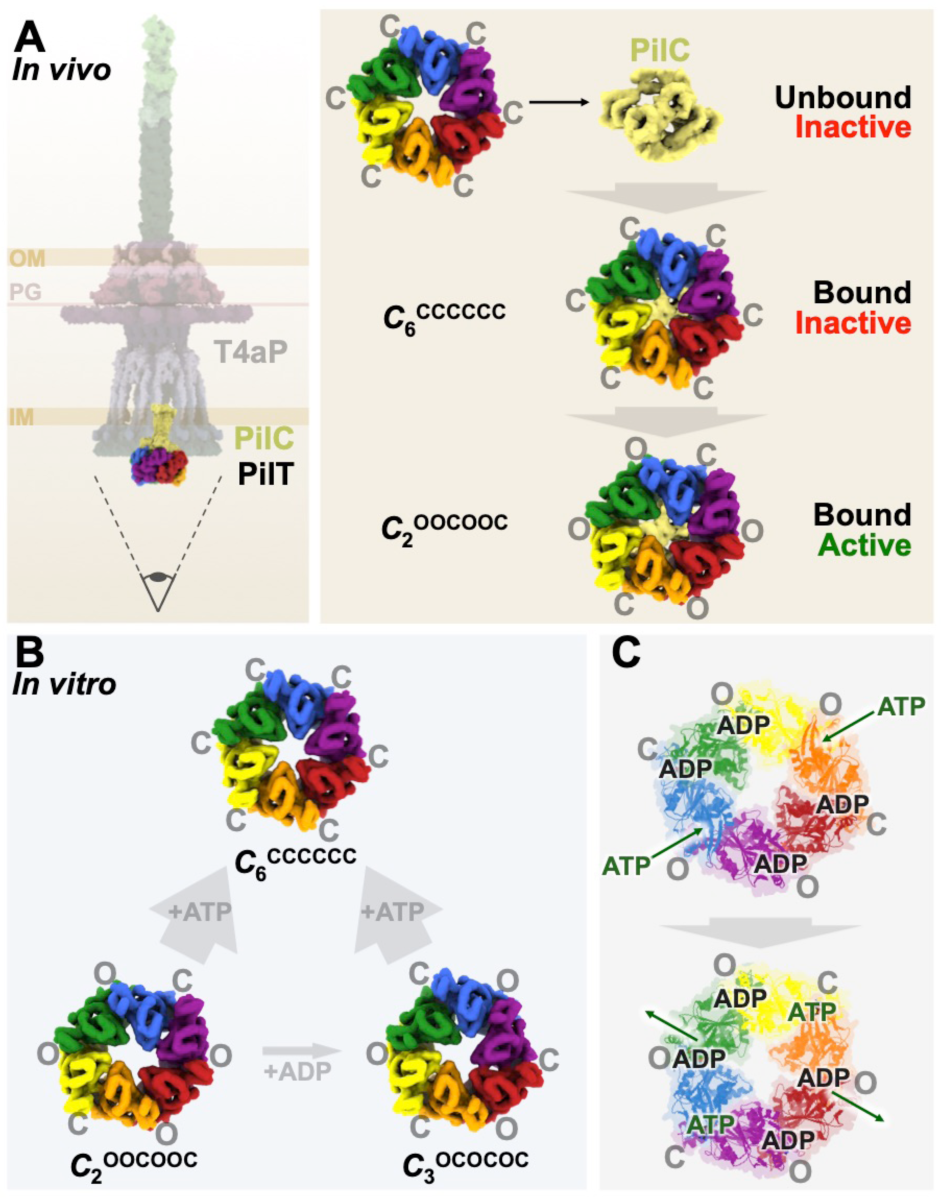
Proposed models of function. **A**, *In vivo* model of PilT conformations. The entire T4aP is shown with an eye symbol (left) indicating the viewing direction for the model (right). This model suggests that the *C*_6_^CCCCCC^ and *C*_2_^OOCOOC^ conformations are of *in vivo* functional relevance. **B**, *In vitro* model of PilT conformations. This model illustrates that *in vitro* PilT is a mixture of *C*_2_^OOCOOC^ and *C*_3_^OCOCOC^ conformations; added ADP can modestly increase the proportion of the *C*_3_^OCOCOC^ conformation while added ATP strongly favours the *C*_6_^CCCCCC^ conformation. **C**, Updated model of ATP binding and hydrolysis in PilT. The O- interfaces of PilT have a higher affinity for ADP than those of PilB, suggesting a model in which ADP remains bound to the O-interfaces of PilT for one cycle of ATP binding and hydrolysis, i.e. at least four nucleotides are bound to PilT at any given moment in the cycle. This would leave only two apo-interfaces available for binding ATP, promoting a single direction of rotation.

Our mutational and coevolution analyses support the *in vivo* importance of the *C*_6_^CCCCCC^ conformation. Based on BACTH data, the *C*_*6*_^CCCCCC^-targeting R123D mutation impairs PilC interaction. Given this analysis supporting its importance, it was initially surprising that the C-interface destabilizing D31K mutation and the O-interface stabilizing K58A mutation promote hyper-twitching. These results suggest that the active motor is likely to contain O-interfaces. We propose that after binding to PilC in the *C*_6_^CCCCCC^ conformation, PilT converts to an O-interface-containing conformation to power pilus depolymerization (**Figure 6A**). Our data suggest that the only O-interface-containing PilT conformations found in solution are *C*_*2*_^OOCOOC^ and *C*_*3*_^OCOCOC^. PilC is proposed to be a dimer *in vivo* (15) and PilC-like proteins have crystallized with *C*_2_ and asymmetric-pseudo-*C*_2_ symmetry (46, 47); thus, when PilT binds PilC, we anticipate the interaction would induce *C*_2_ symmetry on PilT. Since the *C*_*2*_^OOCOOC^ conformation of PilT is the only *C*_2_ symmetric conformation of PilT found in solution, we propose that the active motor O- interface containing conformation is the *C*_*2*_^OOCOOC^ conformation. Adopting other conformations in the presence of ATP may be a strategy to limit unnecessary hydrolysis in the absence of other T4aP proteins, and could explain why the activity of PilT-like ATPases is notoriously low *in vitro* (11, 39).

In our previously proposed *C*_*2*_^OOCOOC^ model of PilT retraction, we suggested that binding of 2 ATPs to opposite O-interfaces caused the interfaces to close (18). This is consistent with our observations that bound ATP correlates with the C-interface. Closure of two interfaces was predicted to open the neighbouring C-interfaces to enable release of ADP (18). Our data herein suggest that ADP may not be released immediately. Unlike in PilB^Gm^ structures (18), ADP was found in both the O- and C-interfaces in PilT^Gm^ structures. This affinity for ADP to the O- interface is consistent with a model in which PilT temporarily retains ADP for an additional round of ATP hydrolysis after the C-interface opens (**Figure 6C**). Such a mechanism would parallel that of PilB (18) despite their different patterns of O- and C-interfaces. The consequence of this nuance would be that for PilB and PilT, only two O-interfaces would be available at any one time for binding ATP. This scenario would commit PilT to a single direction of ATP binding and hydrolysis and thus a single direction of pore rotation. Consistent with the distinction between *C*_*2*_^CCOCCO^ and *C*_*2*_^OOCOOC^ conformations promoting PilB-like and PilT-like functions, respectively, we note that while PilB adopted the *C*_*2*_^CCOCCO^ conformation in solution here and elsewhere (27), the equivalent *C*_2_ symmetric conformation adopted by PilT^Gm^ and PilT^Pa^ is the *C*_*2*_^OOCOOC^ conformation. Thus, the preferred conformations of PilB and PilT in solution correlate with function. Given that neither PilT^Gm^ nor PilT^Pa^ crystallized in the *C*_*2*_^OOCOOC^ conformation, we believe that individual PilT-like ATPase crystal structures should be interpreted with caution in the absence of accompanying cryoEM analysis.

In addition to the *C*_*2*_^OOCOOC^ conformation, PilT is also found in the *C*_*3*_^OCOCOC^ conformation in solution. The function of the *C*_*3*_^OCOCOC^ conformation remains unclear. That a motor capable of rotating a substrate protein would switch between *C*_2_ and *C*_3_ symmetries during its catalytic cycle is unprecedented. As judged by the similarity between O- and C-interfaces between conformations, it may have been prohibitively difficult during evolution to stabilize the *C*_*2*_^OOCOOC^ conformation without also stabilizing other conformations. Perhaps, evolution favoured the relative stability of the O- vs C-interfaces rather than particular hexamer conformations. Thus, the difference between a retraction ATPase and an extension ATPase may be the relative stabilities of their O- and C-interfaces.

Although the C_2_^CCOCCO^ PilT conformation was not observed in solution, finding that a single PilT otholog can adopt both C_2_^CCOCCO^ and C_2_^OOCOOC^ conformations may be critical for understanding PilT-like ATPase function and evolution. This finding suggests that PilT^Gm^ and potentially other PilT-like ATPases have the capacity to switch between *C*_2_^OOCOOC^ powered counter-clockwise pore rotation (i.e. pilin polymerization), and *C*_2_^CCOCCO^ powered clockwise pore rotation (i.e. pilin depolymerization), blurring the line between extension and retraction ATPases. Indeed, PilT is inexplicably essential for T4aP pilin polymerization in *Francisella tularensis* (48). Similarly, some T4P-like systems – including the T2S, T4bP, Tad/Flp pilus, and even some T4aP systems – can retract in the absence of a dedicated retraction ATPase or in PilT-deleted backgrounds (49-51). A similar conformation switch could also explain how FlaI switches between clockwise and counter-clockwise archaellum rotation (52). Such a switch might easily be regulated by post-translational modifications or partner-protein interactions that modulate the relative stability of the O- vs C-interfaces. Indeed, evidence emerged during the completion of this manuscript that the single PilT-like ATPase from the *Caulobacter* Tad pilus system powers both pilus polymerization and depolymerisation (53). It may be that the last common PilT-like ATPase ancestor catalyzed both clockwise and counter-clockwise rotation, facilitating both pilus polymerization and depolymerization, and only more recently have PilB and PilT specialized to perform just one of these functions.

## EXPERIMENTAL PROCEDURES

### Cloning and mutagenesis

PilT4 from *G. metalloreducens* GS-15 was PCR amplified from genomic DNA using primers P168 and P169 (**Table 2**), digested with NdeI and XhoI, and cloned into pET28a with a thrombin-cleavable hexahistadine tag to create pET28a:PilT^Gm^. PilT from *P. aeruginosa* PAO1 was PCR amplified from genomic DNA using primers P220 and P221, digested with KpnI and XbaI, and cloned into pBADGr for arabinose inducible expression to create pBADGr:PilT^Pa^. pKT25::PilT^Pa^ and pBADGr:PilT^Pa^ were mutated by site directed mutagenesis (Quikchange II, Agilent) to generate E204A, H229A, R239A, E219K, D31K, E258A, R294E, D242K, T216R, R123D, K58A, K58Q, and K58R mutants using the eponymously named primers (**Table 2**). Plasmid sequences were verified by TCAG sequencing facilities (The Hospital for Sick Children, Canada).

**Table 2:**
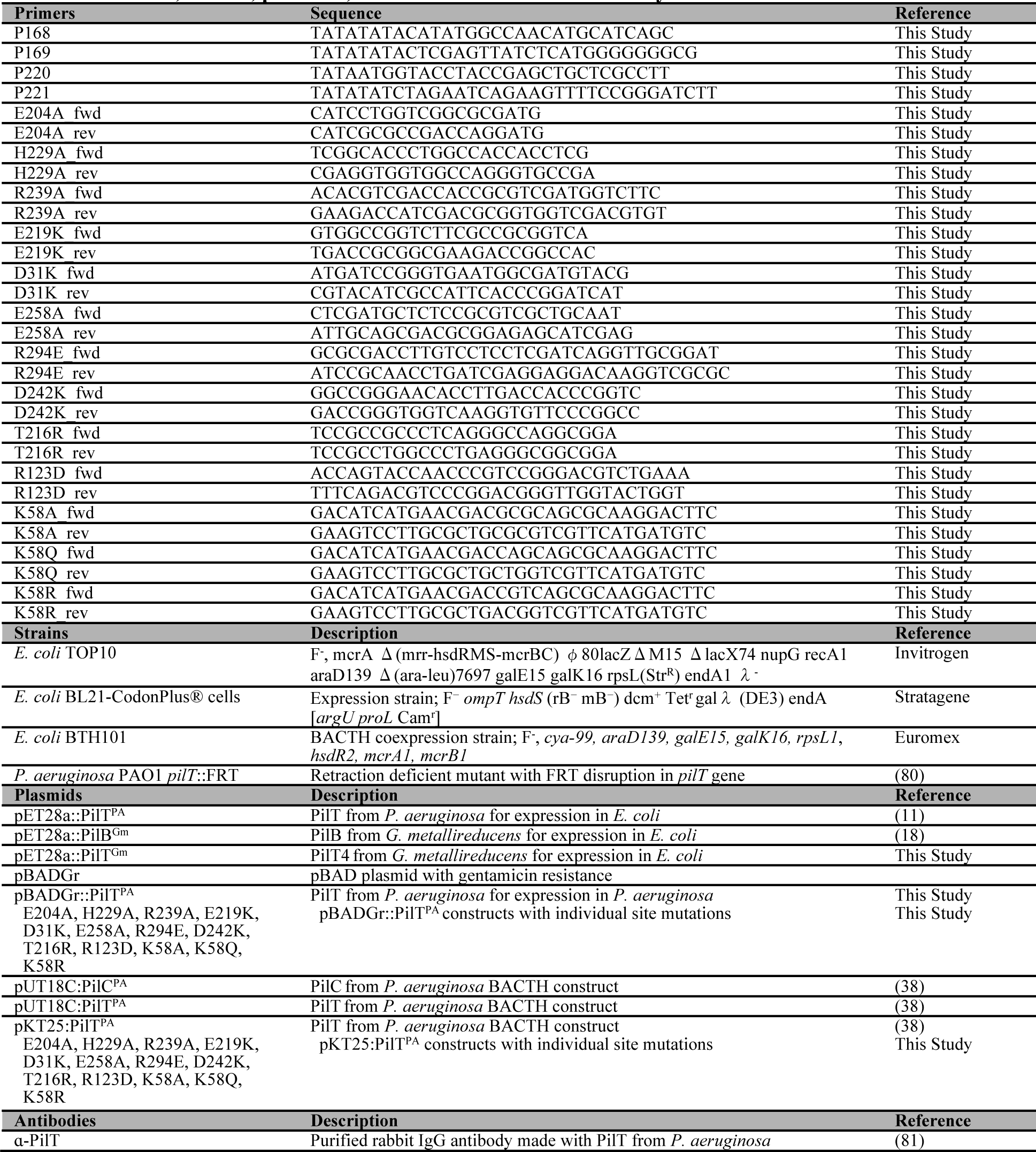
Primers, strains, plasmids, and antibodies used in this study.

### Expression and Purification

*E. coli* BL21-CodonPlus^®^ cells (**Table 2**) were transformed with pET28a:PilT^Gm^, grown in 4 L of lysogeny broth (LB) with 100 µg/ml kanamycin at 37 °C to an A_600_ of 0.5–0.6, then protein expression was induced by the addition of isopropyl-D-1-thiogalactopyranoside (IPTG) to a final concentration of 1 mM, and the cells grown for 16 h at 18 °C. Cells were pelleted by centrifugation at 9000 × *g* for 15 min. Cell pellets were subsequently resuspended in 40 ml binding buffer (50 mM Tris-HCl pH 7, 150 mM NaCl, and 15 mM imidazole). Subsequent to crystallization of methylated *C*_*3*_^OCOCOC^ PilT, the buffer was optimized to improve protein thermostability – the pH was increased, HEPES was used instead of Tris, the concentration of NaCl was increased, and glycerol was added; hereafter the binding buffer was 50 mM HEPES pH 8, 200 mM NaCl, 10% (v/v) glycerol, 15 mM imidazole. After resuspension in binding buffer, the cells were lysed by passage through an Emulsiflex-c3 high-pressure homogenizer, and the cell debris removed by centrifugation for 45 min at 40000 × g. The resulting supernatant was passed over a column containing 5 ml of pre-equilibrated Ni-NTA agarose resin (Life Technologies, USA). The resin was washed with 10 column volumes (CV) of binding buffer and eluted over a gradient of binding buffer to binding buffer plus 600 mM imidazole. Purified protein was next separated on a HP anion exchange column pre-equilibrated with binding buffer; the flow-through contained PilT^Gm^. PilT^Pa^ and PilB^Gm^ were purified as described previously (18, 38). PilT^Gm^, PilT^Pa^, or PilB^Gm^ was then further purified by size exclusion chromatography on a HiLoad™ 16/600 Superdex™ 200pg column pre-equilibrated with binding buffer without imidazole or glycerol. For the *C*_*3*_^OCOCOC^ PilT structure with full-occupancy ADP, 2 mM ATP and 2 mM MgCl_2_ were added just prior to size exclusion chromatography. For the crystallization of methylated *C*_*3*_^OCOCOC^ PilT, the size exclusion chromatography buffer was 50 mM HEPES pH 7, 150 mM NaCl, 10 % v/v glycerol, and subsequent to purification PilT^Gm^ was reductively methylated overnight (Reductive Alkylation Kit, Hampton Research), quenched with 100 mM Tris-HCl pH 7, and the size exclusion chromatography step was repeated. All purified proteins were used immediately.

### Crystallization, Data Collection, and Structure Solution

For crystallization, purified PilT^Gm^ was concentrated to 15 mg/ml (4 mg/ml for methylated *C*_*3*_^OCOCOC^ PilT) at 3000 × *g* in an ultrafiltration device (Millipore). Crystallization conditions were screened using the complete MCSG suite (MCSG 1-4) (Microlytic, USA) using a Gryphon LCP robot (Art Robbins Instruments, USA). Crystal conditions were screened and optimized using vapour diffusion at 20 °C with Art Robbins Instruments Intelli-Plates 96-2 Shallow Well (Hampton Research, USA) with 1 µl protein and 1 µl reservoir solution. For the methylated *C*_*3*_^OCOCOC^ PilT structure, the reservoir solution was 200 mM ammonium sulfate, 100 mM Bis-Tris-HCl pH 6.5, and 25 % (w/v) PEG3350. For cryoprotection of the methylated *C*_*3*_^OCOCOC^ PilT crystal, 2 µl of 50 % (w/v) ethylene glycol, and 50 % (v/v) reservoir solution was added to the drop containing the crystal for 10 s prior to vitrification in liquid nitrogen. For the ADP-bound *C*_*3*_^OCOCOC^ PilT structures, the reservoir solution was 11.5 % (w/v) PEG3350, 200 mM L-proline, and 100 mM HEPES – pH 7.1 for the partial occupancy ADP structure or pH 7.6 for the full occupancy ADP structure. For cryoprotection of ADP-bound *C*_*3*_^OCOCOC^ PilT crystals, 2 µl of 7.5 % (w/v) xylitol, 15 % (w/v) sucrose, and 50 % (v/v) reservoir solution was added to the drop containing the crystal for 10 s prior to vitrification in liquid nitrogen. For the *C*_*6*_^CCCCCC^ PilT structure, the protein solution also contained 2 mM ATP and 2 mM MgCl_2_. The reservoir solution was 9 % (w/v) PEG4000, 100 mM MES pH 6, and 200 mM MgCl_2_. For cryoprotection of the *C*_*6*_^CCCCCC^ PilT crystal, 2 µl of 50 % (w/v) ethylene glycol, and 50 % (v/v) reservoir solution was added to the drop containing the crystal for 10 s prior to vitrification in liquid nitrogen. For the *C*_*2*_^CCOCCO^ PilT structure, the protein solution also contained 2 mM ANP, 2 mM MgCl_2_. The reservoir solution was 2 % (w/v) benzamidine-HCl, 100 mM HEPES-HCl pH 7.9, 200 mM ammonium acetate, and 17.5 % (w/v) PEG3350. For cryoprotection of the *C*_*2*_^CCOCCO^ PilT crystal, 2 µl of 15 % (w/v) xylitol, and 50 % (v/v) reservoir solution was added to the drop containing the crystal for 10 s prior to vitrification in liquid nitrogen.

Diffraction data were collected using synchrotron X-ray radiation as noted in **Table 3**. The data were indexed, scaled, and truncated using XDS (54). The *C*_*2*_^CCOCCO^ PilT data were anisotropically truncated and scaled using the Diffraction Anisotropy Server (55). PHENIX-MR (56) was used solve the structures of PilT^Gm^ by molecular replacement with PDB 3JVV pre-processed by the program Chainsaw (57). In every case, the resulting electron density map was of high enough quality to enable building the PilT protein manually in COOT (58). Through iterative rounds of building/remodelling in COOT (58) and refinement in PHENIX-refine (59) the structures were built and refined. Rosetta refinement in PHENIX (60) helped improve models early in the refinement process, while refinement in the PDB-redo webserver (61) helped improve models late in the refinement process. *C*_*6*_^CCCCCC^ PilT were refined with individual B-factors, all other structures were refined with a single B-factor per residue. The occupancy of the nucleotides in the partial-occupancy-ADP *C*_*3*_^OCOCOC^ PilT structure were estimated by PHENIX-refine, restricting all atoms in a nucleotide to a uniform occupancy. Progress of the refinement in all cases was monitored using *R*_free_.

**Table 3.**
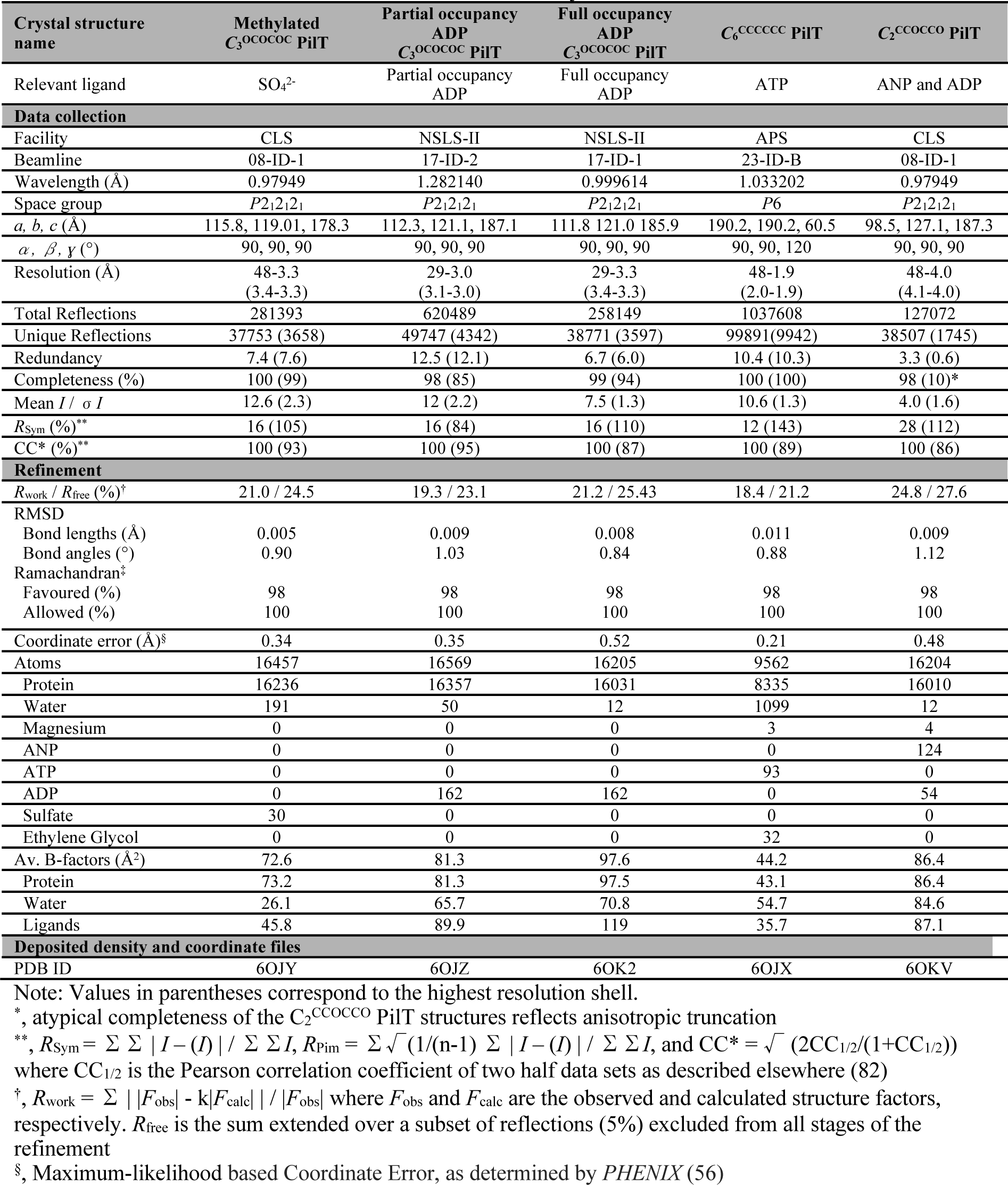
Data collection and refinement statistics of PilT crystal structures.

### Enzyme-coupled ATPase assay

Enzyme-coupled ATPase assays were performed as done elsewhere (62), with minor modifications. Briefly, the reaction buffer included 40 u/ml lactate dehydrogenase (Sigma), 100 u/ml pyruvate kinase (Sigma), 100 mM NaCl, 2 mM MgCl_2_, 25 mM KCl, 0.8 mM nicotinamide adenine dinucleotide, 10 mM phosphoenolpyruvate, 5 mM ATP, and 0.088 mM PilT^Gm^. The pH of the reaction buffer was set with a 200 mM MES and 200 mM HEPES dual buffer, at pH 5.5, 6.0, 6.5, 7.0, 8.5, or 9.0. The reaction volume was 100 µl. Conversion of NADH to NAD+ – proportional to the ADP produced by the hydrolysis of ATP – was monitored by measuring the A_340_ every 2 min at 25°C for 2 h. Initial reaction rates were used. Control experiments were performed (without added PilT^Gm^) spiking the reaction buffer at different pH values with 2 mM ADP; conversion of all NADH to NAD+ occurred almost immediately at every pH value used herein indicating that the reagents were not rate limiting.

### Identifying hexamer symmetry

Each hexamer (extracted from the corresponding PDB coordinates) was aligned in the PyMOL Molecular Graphics System, version 2.2 (Schrodinger, LLC. 2010) against the same hexamer six times. Specifically, all six chains were aligned with the n+1 chains, n+2 chains, n+3 chains, n+4 chains, n+5 chains, or n+6 chains, and the RMSD^C*α*^ of the alignments noted. If the RMSD^C*α*^ was below 1 Å the rotation was considered to be equivalent, and if the RMSD^C*α*^ was above 4 Å the rotation was considered to be distinct; this was used to define the symmetry of the hexamer. For example, in the *C*_*2*_^CCOCCO^ PilT structure the RMSD^C*α*^, the n+3 and n+6 alignments were equivalent while the n+1, n+2, n+4, and n+5 alignments were distinct, consistent with *C*^2^ rotational symmetry. If the RMSD^C*α*^ was between 1 Å and 4 Å, the rotation was considered to be pseudo-symmetric. For example, in the ADP-bound *C*_*3*_^OCOCOC^ PilT structures, the n+1, n+3, n+5 alignments were distinct while the RMSD^C*α*^ of the n+2, n+4, and n+6 alignments was ∼3.5 Å – a pattern consistent with pseudo-*C*_*3*_ symmetry.

### Plotting the residues at the packing unit interface

CMview (63) was used to identify residues that contact one another. The contact type was set to ‘ALL’ (i.e. every atom available in the structure) and the distance cut-off was set to 4 Å. To identify residues that contact one another even in low-resolution structures, where side-chain modeling is less definitive, additional residue contacts were identified by setting the contact type to ‘BB’ (i.e. backbone atoms) and the distance cut-off to 8 Å. To identify tertiary structure contacts, the N2D (residues 1-100 in PilT^Gm^) and CTD (residues 101-353 in PilT^Gm^) were loaded separately – to simplify analysis, the linker between the N2D and CTD was considered to be part of the CTD. To identify packing unit-forming contacts, the N2D^n^ and CTD^n+1^ from adjacent chains were loaded together, and the previously identified tertiary structure contacts were subtracted from this contact list. To identify contacts that form the interface between two adjacent packing units, two adjacent packing units were loaded together, and then the previously identified tertiary structure and packing unit-forming contacts were subtracted from this contact list.

### Identifying O- vs C- packing unit interfaces

The O- and C-interfaces of PilT^Gm^ crystal structures were initially identified by qualitative comparison with the characterized O- and C-interfaces of PilB^Gm^. Subsequent to our finding that the O- and C-interfaces are correlated with intermolecular T221–H230 and T221– L269 distances, the *α*-carbon distances between these residues were measured in PilT^Gm^. In other PilT-like ATPases, the *α*-carbon distances between the residues that correspond with PilT^Gm^ residues T221, H230, or L269 were used. If the T221–H230 distance was greater than 11 Å and the T221–L269 distance was less 11 Å, this interface was classified as O. If the T221–H230 distance was less than 9 Å and the T221–L269 distance was more than 12 Å, this interface classified as C. Interfaces that did not meet these criteria were classified as X-interfaces.

### Identifying evolutionarily-coupled residues

The PilT^Pa^ amino acid sequence was analyzed using the *EVcouplings* webserver (34) with default parameters. Homologs were identified, aligned, and analyzed – 30421 in total. This analysis did not include the last 60 C-terminal residues of PilT as the alignment had too many gaps in this region and default parameters enforce 30 % maximum gaps allowed. Relaxing this parameter to 50 and 75 % maximum gaps allowed more C-terminal residues to be included in the analysis, though some of the evolutionarily-coupled residue pairs identified with default parameters were not discovered. To compensate, we merged the evolutionarily-coupled residues identified with default parameters, with 50 % maximum gaps, and 75 % maximum gaps. This analysis yielded overall coverage from residue 18 to 347, though coupled residues in the last 60 C-terminal residues likely have a lower likelihood of being identified. Only residue pairs with a PLM score greater than 0.2 were included in subsequent analysis. To better understand the significance of these evolutionarily coupled residues, they were compared to the tertiary structure contacts, packing unit contacts, and O- and C-interface contacts identified in CMview.

### CryoEM analysis

Newly purified PilT^Gm^ at 0.5 mg/ml in binding buffer without imidazole or glycerol was incubated with and without 1 mM MgCl_2_ plus 1 mM ATP, 1 mM MgCl_2_ plus 0.1 mM ADP, 1 mM MgCl_2_ plus 1 mM ADP, 1 mM MgCl_2_ plus 1mM ATP and 0.1 mM ADP, or 1 mM MgCl_2_ plus 1mM ATP and 1 mM ADP at 4°C for 10 min before preparing cryo-EM grids. PilT^Pa^ at 0.75 mg/ml or PilB^Gm^ at 0.6 mg/ml in binding buffer without imidazole or glycerol were also incubated without added nucleotide at 4°C for 10 min before preparing cryo-EM grids. Three µl of protein sample was applied to nanofabricated holey gold grids (64-66), with a hole size of ∼1 µm and blotted using a modified FEI Vitribot Mark III at 100% humidity and 4 °C for 5.5 s before plunge freezing in a 1:1 mixture of liquid ethane and liquid propane held at liquid nitrogen temperature (67).

Cryo-EM data was collected at the Toronto High-Resolution High-Throughput cryo-EM facility. Micrographs from untilted specimens were acquired as movies with a FEI Tecnai F20 electron microscope operating at 200 kV and equipped with a Gatan K2 Summit direct detector camera. Movies, consisting of 30 frames at two frames per second, were collected with defocuses ranging from 1.2 to 3.0 µm. Data were recorded with an exposure rate of 5 electrons/pixel/s with a calibrated pixel size of 1.45 Å/pixel. For the 40° tilted data collection, micrographs were acquired as movies with a FEI Titan Krios electron microscope (Thermo Fisher Scientific) operating at 300 kV and equipped with a Falcon 3EC direct detector camera. Movies, consisting of 30 frames at two seconds per frame, were collected with defocuses ranging from 1.7 to 2.5 µm. Data were recorded with an exposure rate of 0.8 electrons/pixel/s with a calibrated pixel size of 1.06 Å/pixel.

All image processing of the cryoEM data was performed in *cryoSPARC v2.8.0* (36) (**Table 4**). Movie frames were aligned with an implementation of *alignframes_lmbfgs* within *cryoSPARC v2* (68) and CTF parameters were estimated from the average of aligned frames with *CTFFIND4* (69). Initial 2D class averages were generated with manually selected particles; these classes were then used to select particles. Particle images were selected and beam-induced motion of individual particles corrected with an improved implementation of *alignparts_lmbfgs* within *cryoSPARC v2* (68). For the 40° tilted data, an implementation of *GCTF* (37) wrapped within *cryoSPARC v2* was used to refine the micrograph CTF parameters while also locally refining the defocus for individual particles with default parameters (local_radius of 1024, local_avetype set to Gaussian, local_boxsize of 512, local_overlap of 0.5, local_resL of 15, local_resH of 5, and refine_local_astm set to Z-height). Particle images were extracted in 256 × 256 pixel boxes. Candidate particle images were then subjected to 2D classification. For the particles that preferentially adopted top-views, comparison of 2D class averages of these top-views with 2D projections of PilT structures was used to identify the corresponding conformation. 2D projections of PilT structures were generated using *genproj_fspace_v1_01* (J. Rubinstein, https://sites.google.com/site/rubinsteingroup/3-d-fourier-space). For the purposes of 3D classification, particle images contributing to 2D classes without high-resolution features were removed. For samples with tilted-views, *ab initio* reconstruction was performed using 2 to 4 classes. *Ab initio* classes consistent with hexamers were used as initial models for heterogeneous refinement; particles from 3D classes that did not converge at this stage were removed. Particles from distinct 3D classes were then subjected to homogeneous refinement.

**Table 4.**
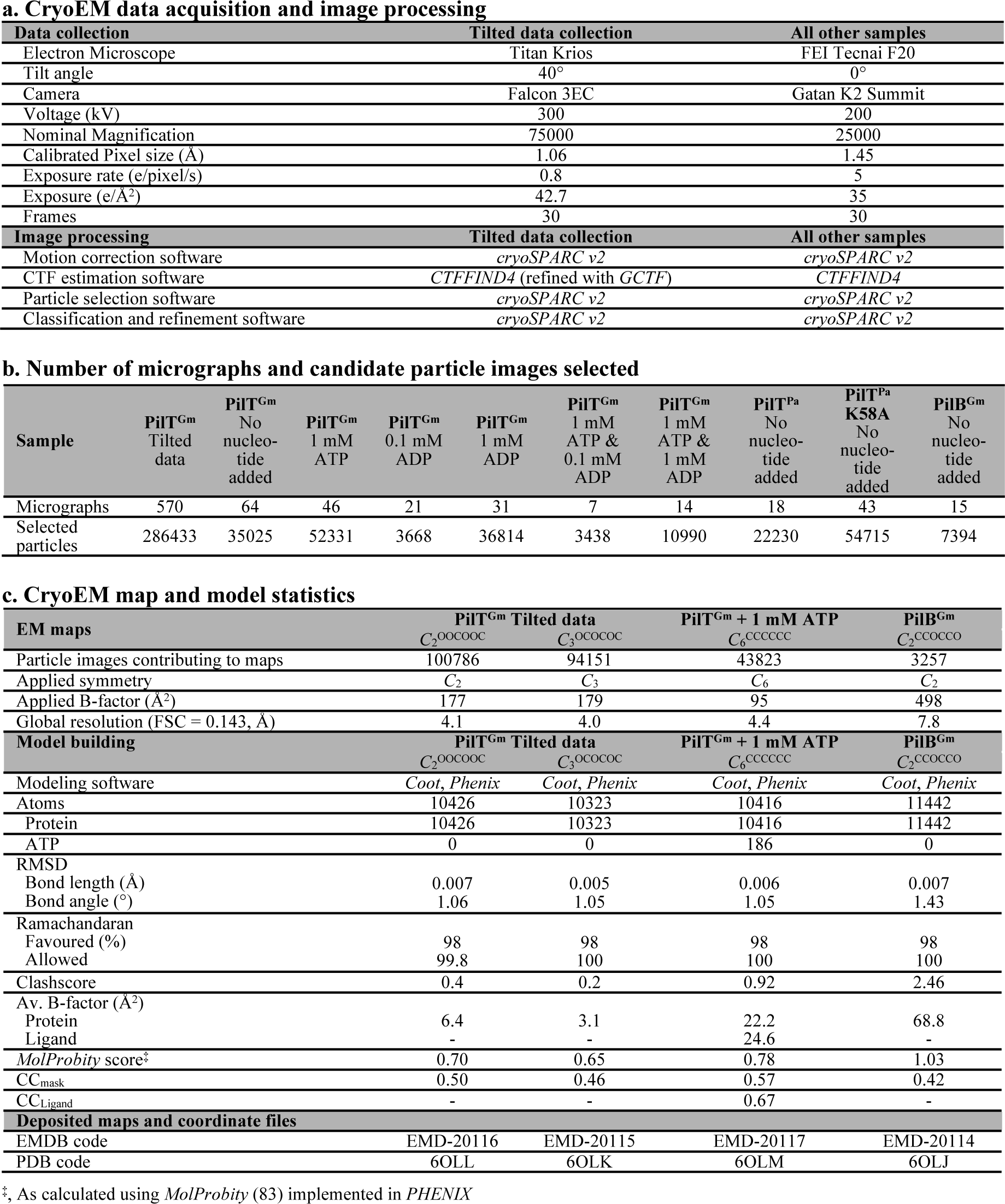
CryoEM data acquisition, processing, atomic model statistics and map/model depositions.

Molecular models could be built into these maps by fitting rigid packing units of the PilT^Gm^ crystal structures into the maps in Chimera (70). These models were refined against the maps in Phenix-Refine (71) with reference model restraints to the ATP bound PilT^Gm^ crystal structure (or for the PilB^Gm^ model, PDB 5TSH), enabling the following refinement options: minimization_global, rigid_body, simulated_annealing, and adp. Side-chains were not modeled.

### *In vivo* twitching motility assays, PO4 phage infection, sheared surface pilin preparation, and western blotting

*P. aeruginosa* PAO1 *pilT*::FRT (**Table 2**) was electroporated with pBADGr, pBADGr::PilT^Pa^, or pBADGr::PilT^Pa^ derivative mutant constructs for transcomplementation of PilT. Twitching assays were performed as described previously (72, 73) except that assays were performed in 150 mm by 15 mm petri polystyrene dishes (Fisher Scientific) with 30 µg/ml gentamicin for 18 h at 37 °C. After this incubation, the agar was then carefully discarded and the adherent bacteria was stained with 1 % (w/v) crystal violet dye, followed by washing with deionized water to remove unbound dye. Twitching zone areas were measured using ImageJ software (74). Twitching motility assays were performed in six replicates.

Surface pili were analyzed as previously described (75), with the exception that sheared supernatant pili and flagellins were precipitated with 100 mM MgCl_2_ incubated at room temperature for 2 h prior to pelleting. These pellets were resuspended in 100 *μ*l of 1× SDS-PAGE sample buffer, 10 *μ*l of which was loaded onto a 20 % SDS-PAGE gel. In parallel, western blot analysis was performed on the cells used to produce the pili to confirm mutant stability using rabbit polyclonal anti-PilT antibodies (**Table 2**).

In preparation for the PO4 phage infection assay, 25 ml LB with 1.5 % (w/v) agar was solidified in 150 mm by 15 mm polystyrene petri dishes (Fisher Scientific). On top of this layer, 8 ml of 0.6 % (w/v) agar pre-inoculated with 1 ml of A_600_ = 0.6 transcomplemented *P. aeruginosa* PAO1 *pilT*::FRT was poured and solidified. Three µl of 10^4^ plaque forming units per ml PO4 phage was then spotted onto these plates in triplicate and incubated at 30°C for 16 h before images of the plates were acquired.

### Bacterial Two-hybrid analysis (BACTH)

BACTH analysis was performed as done previously (38) with minor adjustments. Briefly, *E. coli* BTH101 cells (**Table 2**) were co-transformed with pUT18C::pilC, or pUT18C::pilT and mutants of pKT25::pilT. Preliminary tests suggested that prolonged incubation in induction conditions at 25 °C improved the signal to noise, so three colonies from each of these transformations were individually streaked onto MacConkey-Maltose Agar plus 100 µM Ampicillin, 50 µM Kanamycin, and 0.5 mM IPTG and incubated over-night at 25 °C. A single colony from these plates were then used to inoculate 500 µl of LB plus 100 µM Ampicillin, 50 µM Kanamycin, and 0.5 mM IPTG at 25 °C until the A_600_ was 1.0. 5 µl of this solution was then used to inoculate 500 µl of LB plus 100 µM Ampicillin, 50 µM Kanamycin, and 0.5 mM IPTG at 25°C for 16 hours. 35 µl of this was then used to inoculate 600 µl of LB plus 100 µM Ampicillin, 50 µM Kanamycin, and 0.5 mM IPTG at 25 °C for 1 hour, then at 18 °C until the A_600_ = 0.6. All the replicates were then normalized to 600 µl and A_600_ = 0.6, then pelleted at 3200 × *g* for 5 minutes. The supernatant was carefully removed, and the pellet was resuspended in 100 µl 200 mM Na_2_HPO_4_ pH 7.4, 20 mM KCl, 2 mM MgCl_2_, 0.8 mg/ml cetyltrimethylammonium bromide detergent, 0.4 mg/ml sodium deoxycholate, and 0.54 % (v/v) *β*-mercaptoethanol. After 5 minutes of incubation at room temperature, 10 µl of this solution was then transferred to a 96 well clear bottom plate. 150 µl of a second solution was then added: 60 mM Na_2_HPO_4_ pH 7.4, 40 mM KCl, 20 µg/ml cetyltrimethylammonium bromide detergent, 10 µg/ml sodium deoxycholate, 0.27 % (v/v) *β*-mercaptoethanol, and 1 mM *ortho*-nitrophenyl-*β*-galactoside. The A_420_ and A_550_ were measured every 2 minutes for 30 minutes at 30 °C. *β*-galactosidase activity in Miller Units was calculated by finding the slope of 1000 × (A_420_ - 1.75 × A_550_) / (0.6 absorbance units × 0.06 ml) over time (minutes) of the linear portion of the initial reaction.

## Acknowledgements

We thank Roland Pfoh, Jeff Lee, and Natalie C. Bamford, as well as Zev Ripstein, Hui Guo, and Thamiya Vasanthakumar for technical assistance during crystal diffraction screening/data collection and cryoEM grid freezing/data collection, respectively. Shane Caldwell is thanked for technical assistance and helpful discussions. This work was supported by a grant from the Canadian Institutes for Health Research (CIHR) MOP 93585 to L.L.B. and P.L.H. M.M. was supported by a CIHR Doctoral Studentship and Ontario Graduate Scholarship during these studies. P.L.H. and J.L.R. are recipients of Canada Research Chairs. Funds for the X-ray and EM facilities at The Hospital for Sick Children were provided in part by the Canadian Foundation for Innovation and the Government of Ontario. Protein crystals were grown and screened in the Structural & Biophysical Core (SBC) Facility at The Hospital for Sick Children. The crystallographic and graphics programs used in this study were accessed using SBGrid.

---

**Figure 1 – figure supplement 1.**
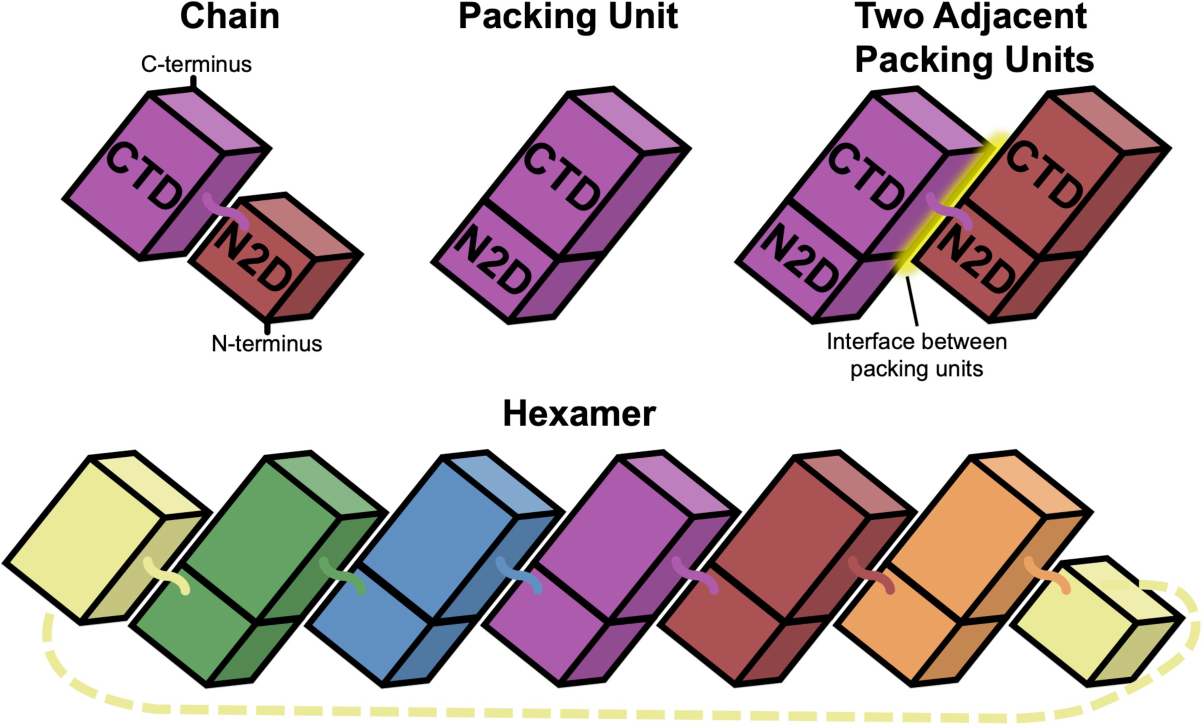
Nomenclature used herein. Each PilT chain is composed of two domains: N2D and CTD. The packing unit is composed of the NTD^n^ and CTD^n+1^. Two adjacent packing units contact one-another creating an interface between packing units. Six packing units organize in this manner to create a hexamer.

**Figure 4 – figure supplement 1.**
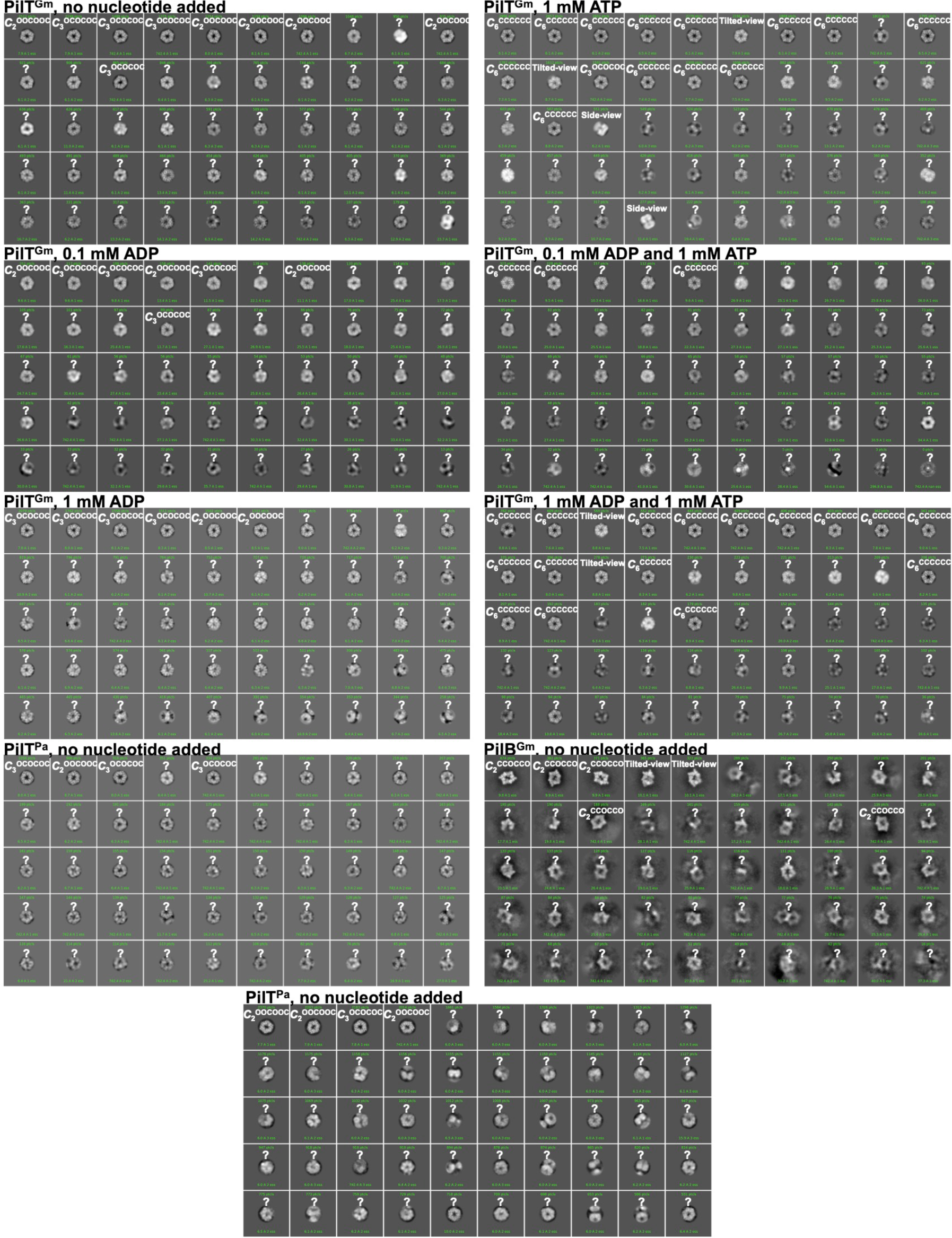
2D class averages of PilB and PilT particles and their proposed conformation. The protein and nucleotide added are indicated above each list of 50 class averages. Class averages are sorted by the number of particles in each class. The qualitative assignment of the class average conformation is noted above each class average in white. White question marks indicate class averages that were too blurry to identify the conformation. Class averages that appear to represent tilted-views or side views are also labeled in white.

**Figure 4 – figure supplement 2.**
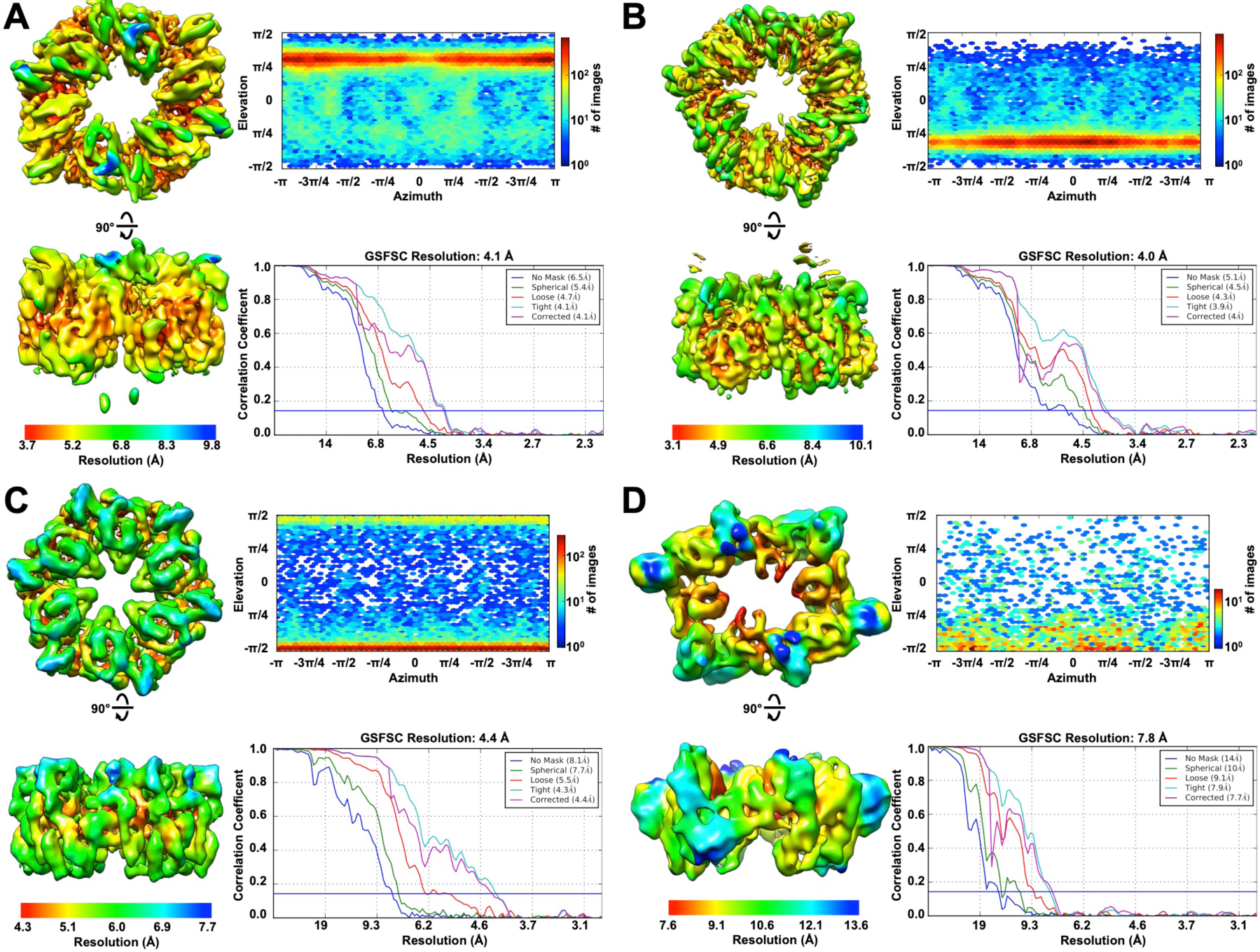
Local resolution (left), particle distribution (upper right), and Fourier shell correlation (FSC) curves (lower right). **A**, PilT^Gm^ *C*_2_^OOCOOC^ conformation map. **B**, PilT^Gm^ *C*_3_^OCOCOC^ conformation map. **C**, PilT^Gm^ *C*_6_^CCCCCC^ conformation map. **D**, PilB^Gm^ *C*_2_^CCOCCO^ conformation map.

